# Endocrine signaling mediates asymmetric motor deficits after unilateral brain injury

**DOI:** 10.1101/2020.04.23.056937

**Authors:** Nikolay Lukoyanov, Hiroyuki Watanabe, Liliana S. Carvalho, Olga Nosova, Daniil Sarkisyan, Mengliang Zhang, Marlene Storm Andersen, Elena A. Lukoyanova, Vladimir Galatenko, Alex Tonevitsky, Igor Bazov, Tatiana Iakovleva, Jens Schouenborg, Georgy Bakalkin

**Author notes:** These authors contributed equally to this work (Nikolay Lukoyanov, Hiroyuki Watanabe, Liliana S. Carvalho, Olga Nosova, Daniil Sarkisyan). Co-senior authors (Jens Schouenborg, Georgy Bakalkin).

## Abstract

A paradigm in neurology is that brain injury-induced motor deficits (e.g. hemiparesis and hemiplegia) arise due to aberrant activity of descending neural pathways. We discovered that a unilateral injury of the hindlimb sensorimotor cortex of rats with completely transected thoracic spinal cord produces hindlimb postural asymmetry with contralateral flexion, and asymmetric changes in nociceptive hindlimb withdrawal reflexes and gene expression patterns in lumbar spinal cord. The injury-induced postural effects were abolished by prior hypophysectomy and were mimicked by transfusion of serum from animals with unilateral brain injury. Antagonists of the opioid and vasopressin receptors blocked formation of hindlimb postural asymmetry suggesting that these neurohormones mediate effects of brain injury on lateralized motor responses. Our data indicate that descending neural control of spinal circuits is complemented by a previously unknown humoral signaling from injured brain to the contra- and ipsilesional hindlimbs, and suggest the existence of a body side-specific neuroendocrine regulation in bilaterally symmetric animals.

## Introduction

Motor deficits secondary to stroke and traumatic brain injury (TBI) are characterized by paralysis (hemiplegia) or weakness (hemiparesis) that are typically developed on the contralesional side of the body (Fernandes et al., 2018; Lemon, 2008; Purves et al., 2001; Roelofs et al., 2018; Smith et al., 2017; Ward, 2017; Wilson et al., 2017). Flaccid paralysis is replaced by spasticity and hyperreflexia. The patients often demonstrate hyperactive stretch reflexes, clonus, clasp-knife response and positive Babinski signs that differ between the left and right extremities. Balance is disturbed with impairments in symmetry, steadiness and dynamic stability (Lemon, 2008; Purves et al., 2001; Roelofs et al., 2018; Smith et al., 2017; Ward, 2017; Wilson et al., 2017). The survivors typically have asymmetry with most of the weight shifted toward the stronger side when in sitting or standing position, and poor postural responses in quiet and perturbed balance (Fernandes et al., 2018). The current paradigm is that these deficits are developed due to aberrant activity of neural tracts descending from the brain to the spinal cord (Lemon, 2008; Purves et al., 2001; Smith et al., 2017; Ward, 2017; Wilson et al., 2017; Wolpaw, 2012).

Animal studies demonstrate that a unilateral brain injury (UBI) induces the hindlimb postural asymmetry (HL-PA) with ipsi- or contralesional limb flexion, and that the developed asymmetry is retained after transection of the spinal cord (Chamberlain et al., 1963; DiGiorgio, 1929; Wolpaw, 2012). Consistent with this, monosynaptic and polysynaptic hindlimb reflexes are enhanced on the ipsilateral side after lateral hemisection of the spinal cord, and this pattern is sustained after spinal transection (Hultborn & Malmsten, 1983; Rossignol & Frigon, 2011). Thus asymmetric motor dysfunctions after brain or spinal cord injury may develop due to the spinal neuroplastic changes induced by abnormal activity of descending pathways.

In this study we challenge the neurological paradigm by investigating whether a unilaterally injured brain may signal to the lumbar spinal cord through non-spinal mechanism. We applied a “reversed strategy” protocol in which brain injury was performed after complete transection of the spinal cord at a superior thoracic level.

## Results

### Brain injury induces postural asymmetry in rats with transected spinal cord

We first demonstrated that the unilateral injury of the hindlimb representation area of the sensorimotor cortex in rats induces formation of HL-PA (***Figure 1A,D,E***; ***Figure 1—figure supplement 1***). The effect was evident under pentobarbital anesthesia within 5 min after the lesion and lasted for 14 days. The HL-PA median values and probability to develop HL-PA greater than the 1 mm threshold were markedly higher in rats with UBI (n = 8) compared to those with sham surgery (sham; n = 7). The UBI rats displayed a contralesional hindlimb flexion which correlated with motor deficits of the same limb in the beam-working and ladder rung tests (UBI, n = 11/12; sham, n = 8) (***Figure 1B,C***). The UBI rats showed the high number of slips of the contralesional hindlimb compared to the ipsilesional limb, and to both hindlimbs in rats with sham surgery. Consistent with earlier studies (Chamberlain et al., 1963; DiGiorgio, 1929; Hultborn & Malmsten, 1983; Rossignol & Frigon, 2011; Wolpaw, 2012), the HL-PA was retained after complete transection of the spinal cord performed at the T2-T3 level (***Figure 1—figure supplement 1E-G***). We hypothesized that the HL-PA is maintained either due to neuroplastic changes in the lumbar spinal cord induced through the descending neural tracts before spinalization, or due to non-spinal cord mediated signaling from the injured brain to the lumbar neural circuits.

**Figure 1.**
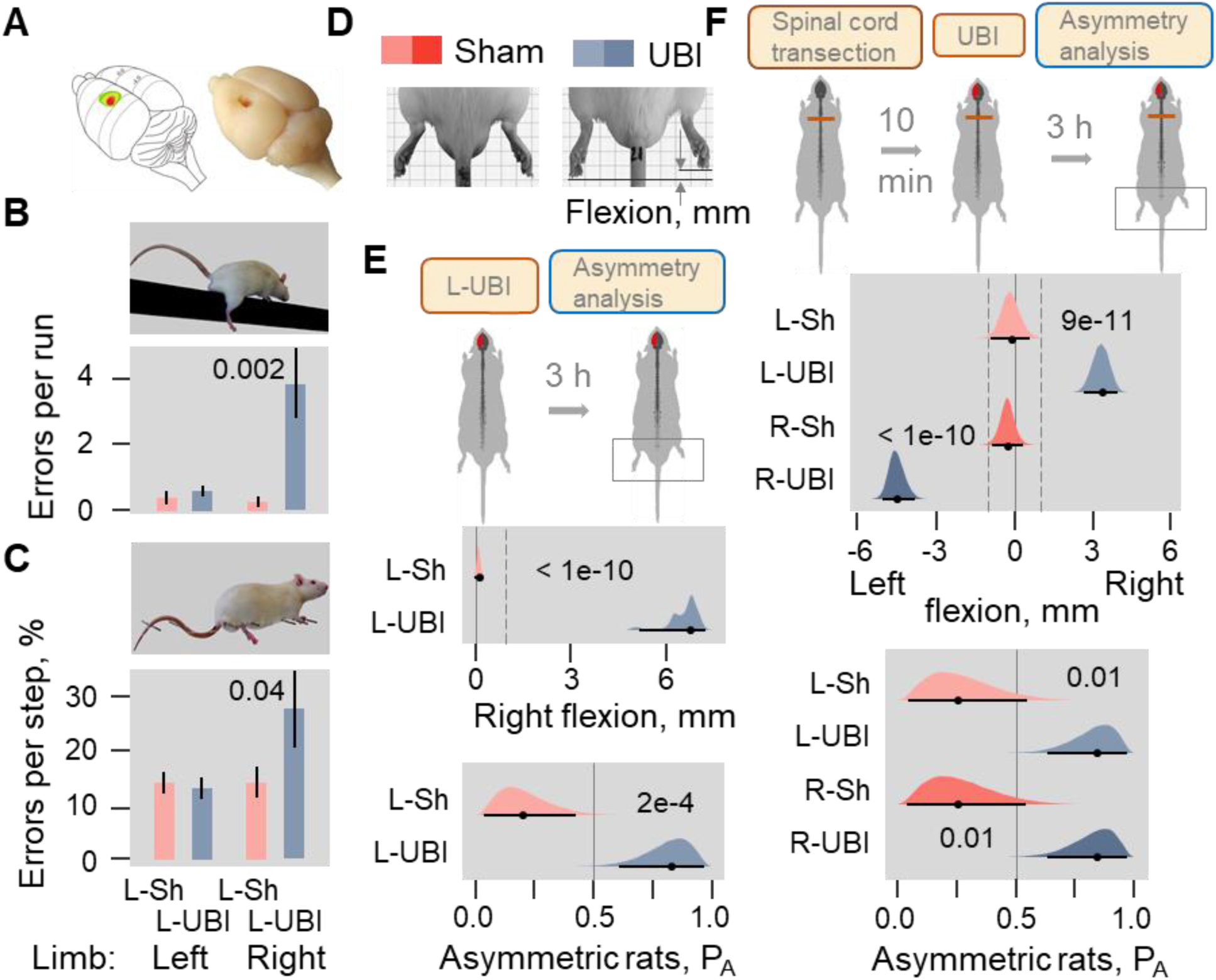
Postural asymmetry of hindlimbs induced by the unilateral ablation of the hindlimb representation area of sensorimotor cortex in rats with intact and completely transected spinal cord. (**A**) Location of the right hindlimb representation area on the rat brain surface (adapted from (Frost et al., 2013)) and a representative UBI. (**B,C)** The left UBI (L-UBI)- produced motor deficits in the beam-working and ladder rung tests. Data are Mean ± SEM analyzed by two-way ANOVA followed by Tukey HSD post-hoc test. (**B**) A main effect of left UBI (F_1,36_ = 8.21, P = 0.007) and hindlimb side (F_1,36_ = 5.80, P = 0.021); interaction: F_1,36_ = 6.78, P = 0.01. Right limb in the left UBI (n = 12) versus either each hindlimb in left sham surgery (L-Sh; n = 8; P = 0.002), or left limb in the left UBI (P = 0.002) group. (**C**) A main effect of the left UBI (F_1,34_ = 3.43, P = 0.072) and hindlimb side (F_1,34_ = 4.11, p = 0.050); interaction: F_1,34_ = 4.22, P = 0.048. Right limb in the left UBI (n = 11) versus either right limb in left sham surgery (n = 8; P = 0.043) or left limb in the left UBI (P = 0.017) group. (**D**) HL-PA analysis. (**E**) HL-PA 3 h after left UBI (n = 8) or left sham surgery (n = 7). (**F**) HL-PA 3 h after left UBI (n = 9) or right UBI (n = 9), and left (n = 4) or right (n = 4) sham surgery, all performed after complete spinal cord transection. In (**E,F**), the HL-PA in millimeters (mm) and probability (P_A_) to develop HL-PA above 1 mm threshold (shown by vertical dotted lines) are plotted as median, 95% HPDC intervals, and posterior distribution from Bayesian regression. Negative and positive HL-PA values are assigned to rats with the left and right hindlimb flexion, respectively. Significant asymmetry and differences between the groups: 95% HPDC intervals did not include zero value, and adjusted P-values were ≤ 0.05. Adjusted P is shown for significant differences identified by Bayesian regression. **Source data 1.** The EXCEL source data file contains data for panels **E** and **F** of ***Figure 1***. **Figure supplement 1.** UBI-induced HL-PA formation and its fixation after complete spinal cord transection. **Figure supplement 2.** Time-course of HL-PA formation after the left and right UBI in rats with transected spinal cord. **Figure supplement 3.** Replication experiments 1 and 2.

We tested the second hypothesis in rats that had complete transection of the spinal cord at the T2- 3 level before the UBI was performed (***Figure 1F***; ***Figure 1—figure supplement 2***; ***Figure 1— figure supplement 3*** showing data of two replication experiments). We observed that within 3 hours following UBI (Left UBI, n = 26; Right UBI, n = 15; sham surgery, n = 29) the rats with transected spinal cord developed HL-PA; the HL-PA values and probability to develop HL-PA were much higher than in rats with sham surgery, and similar to those of the UBI animals with intact spinal cords. Strikingly, the left and right UBI produced HL-PA with contralesional hindlimb flexion that was on the right or left side, respectively. We conclude that HL-PA formation in animals with transected spinal cord is mediated through a non-spinal pathway that decussates the midline and assures the development of contralesional flexion. The HL-PA phenomenon was further analyzed after the left UBI.

### Brain injury induces asymmetry in withdrawal reflexes in rats with transected spinal cord

The nociceptive withdrawal reflexes are instrumental in investigation of pathological changes in hindlimb neural circuits given that the changes are induced by converging inputs from peripheral afferents and descending motor commands (Schouenborg, 2002; Spaich et al., 2014). We next sought to determine whether UBI in rats with transected spinal cord produces differences in nociceptive contra- and ipsilesional hindlimb withdrawal reflexes. Electromyographic responses were recorded from the extensor digitorum longus, interosseous, peroneus longus and semitendinosus muscles of the contra- and ipsilesional hindlimbs in the rats with UBI (n = 18) or sham surgery (n = 11) performed after complete spinal transection (***Figure 2***; ***Figure 2—figure supplement 1***; ***Figure 2—figure supplement 2***). Analysis of the electrically evoked electromyographic responses revealed marked significant differences in the asymmetry index (AI = log_2_[Contra / Ipsi], where Contra and Ipsi were values for muscles of the contra- and ipsilesional limbs) from its zero value in the current threshold for the semitendinosus muscle, and in the number of spikes for the extensor digitorum longus and semitendinosus muscles in UBI rats (***Figure 2C,D***). No asymmetry was developed after sham surgery. Representative UBI-induced asymmetry in the number of spikes for the semitendinosus muscle is shown in ***Figure 2A,B*** (for those of extensor digitorum longus, interosseous and peroneus longus muscles, see ***Figure 2— figure supplement 1***).

**Figure 2.**
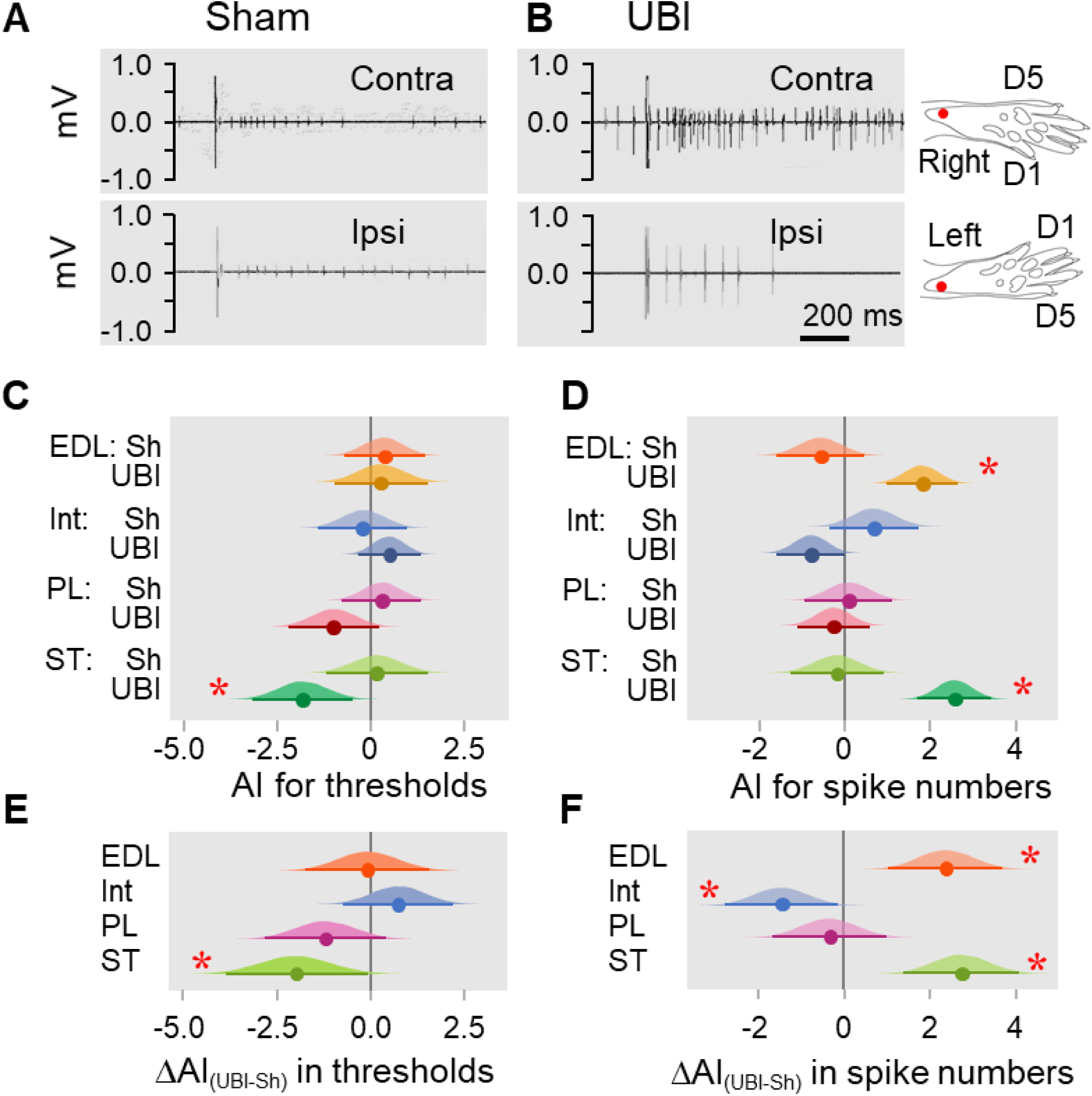
Hindlimb nociceptive withdrawal reflexes in rats exposed to UBI after complete spinal cord transection. EMG activity of left and right extensor digitorum longus (EDL), interosseous (Int), peroneus longus (PL) and semitendinosus (ST) muscles were evoked by electrical stimulation of symmetric paw sites. (**A,B**) Representative semitendinosus responses. (**C,D**) Asymmetry index (AI=log_2_[Contra/Ipsi]) for threshold and spike number. Significance of differences in the asymmetry index from zero value in UBI rats in (**C**) the current threshold for the semitendinosus muscle {median of the posterior distribution (median) = -1.840, 95% highest posterior density continuous interval (HPDCI) = [-3.169, -0.477], adjusted P-value (P) = 0.015, fold difference = 3.6}; and in (**D**) the number of spikes for the extensor digitorum longus (median = 1.818, HPDCI = [0.990, 2.655], P = 4×10^−5^, fold difference = 3.5) and semitendinosus (median = 2.560, HPDCI = [1.691, 3.415], P = 1×10^−8^, fold difference = 5.9) muscles. (**E,F**) Differences in the asymmetry index between the UBI and sham surgery (Sh) groups [ΔAI_(UBI – Sh)_]. Significance of differences in the asymmetry index between the UBI and sham surgery groups for (**E**) the current threshold of the semitendinosus (median = -1.992, HPDCI = [-3.911, -0.106], P = 0.040, fold difference = 4.0); and (**F**) the number of spikes of the interosseous (median = -1.463, HPDCI = [- 2.782, -0.159], P = 0.028, fold difference = 2.8), extensor digitorum longus (median = 2.379, HPDCI = [1.080, 3.743], P = 4×10^−4^, fold difference = 5.2) and semitendinosus (median = 2.745, HPDCI = [1.419, 4.128], P = 6×10^−5^, fold difference = 6.7). Medians, 95% HPDC intervals and densities from Bayesian sampler are plotted. *Significant asymmetry and differences between the groups: 95% HPDC intervals did not include zero value, and adjusted P-values were ≤ 0.05. **Source data 2.** The EXCEL source data file contains data for panels **C** and **D** of ***Figure 2***. **Figure supplement 1.** Effects of the UBI on hindlimb nociceptive withdrawal reflexes. **Figure supplement 2.** Table. The number of rats analyzed in EMG experiments.

The UBI when compared to sham surgery substantially and significantly decreased the asymmetry index for the current threshold of the semitendinosus, and for the number of spikes of the interosseous, while elevated the asymmetry index for the number of spikes of the extensor digitorum longus and semitendinosus (***Figure 2E,F***). No significant changes in peroneus longus were evident. Thus, the nociceptive withdrawal reflexes of the extensor digitorum longus and semitendinosus muscles were asymmetric in the rats with transected spinal cord after the UBI. UBI decreased the threshold for contralesional semitendinosus, and concomitantly activated the contralesional extensor digitorum longus and semitendinosus reflexes and ipsilesional interosseous reflex. This pattern is consistent with formation of the UBI induced contralesional flexion.

### Brain injury produces molecular changes in the lumbar spinal cord

We examined whether the UBI performed after complete spinal transection produces molecular changes in the lumbar spinal segments. Expression of twenty neuroplasticity, opioid and vasopressin genes (***Figure 3—figure supplement 1***), and the levels of three opioid peptides were analyzed in the ipsi- and contralesional halves of the lumbar spinal cord of the rats with transected spinal cord that were exposed to the left UBI (n = 12) or left sham surgery (n = 11). Opioid and vasopressin neurohormones were included because of their involvement in spinal asymmetric processes (see next section). The median expression asymmetry index (eAI = log_2_[Contra/Ipsi], where Contra and Ipsi were the levels in the contra- and ipsilesional lumbar domains) of 19 out of 20 genes at the pairwise comparison was lower in the UBI compared to sham group (sign-test: P = 4×10^−5^) (***Figure 3A,B***). Among these 19 genes, the expression asymmetry index was decreased for *Syt4* (P = 0.004; ***Figure 3C***), and for *Oprk1, Oprm1, Dlg4* and *Homer1* (P _un-adjusted_ < 0.05; ***Figure 3—figure supplement 2A-D***). Changes in the expression asymmetry index were due to elevated expression of 15 genes (sign test, P = 0.041) including *Syt4, Grin2a, Grin2b* and *Oprk1* (for all four, P _un-adjusted_ < 0.05) in the ipsilesional domain (***Figure 3—figure supplement 2E-H***; ***Figure 3—figure supplement 3A-D***) concomitantly with decreased expression of 17 genes (sign test, P = 0.003) including *Gap43* and *Penk* (for both, P _un-adjusted_ < 0.05) in the contralesional domain (***Figure 3—figure supplement 2I,J***; ***Figure 3—figure supplement 3A-D***). The left UBI elevated the levels of opioid peptide Met-enkephalin-Arg-Phe, the proenkephalin marker, in the ipsilesional (P = 9×10^−4^) and contralesional halves (P _un-adjusted_ = 0.020) (***Figure 3D***) and the prodynorphin-derived dynorphin B and Leu-enkephalin-Arg in the ipsilesional domain (for both, P _un-adjusted_ < 0.05) (***Figure 3—figure supplement 2K,L***).

**Figure 3.**
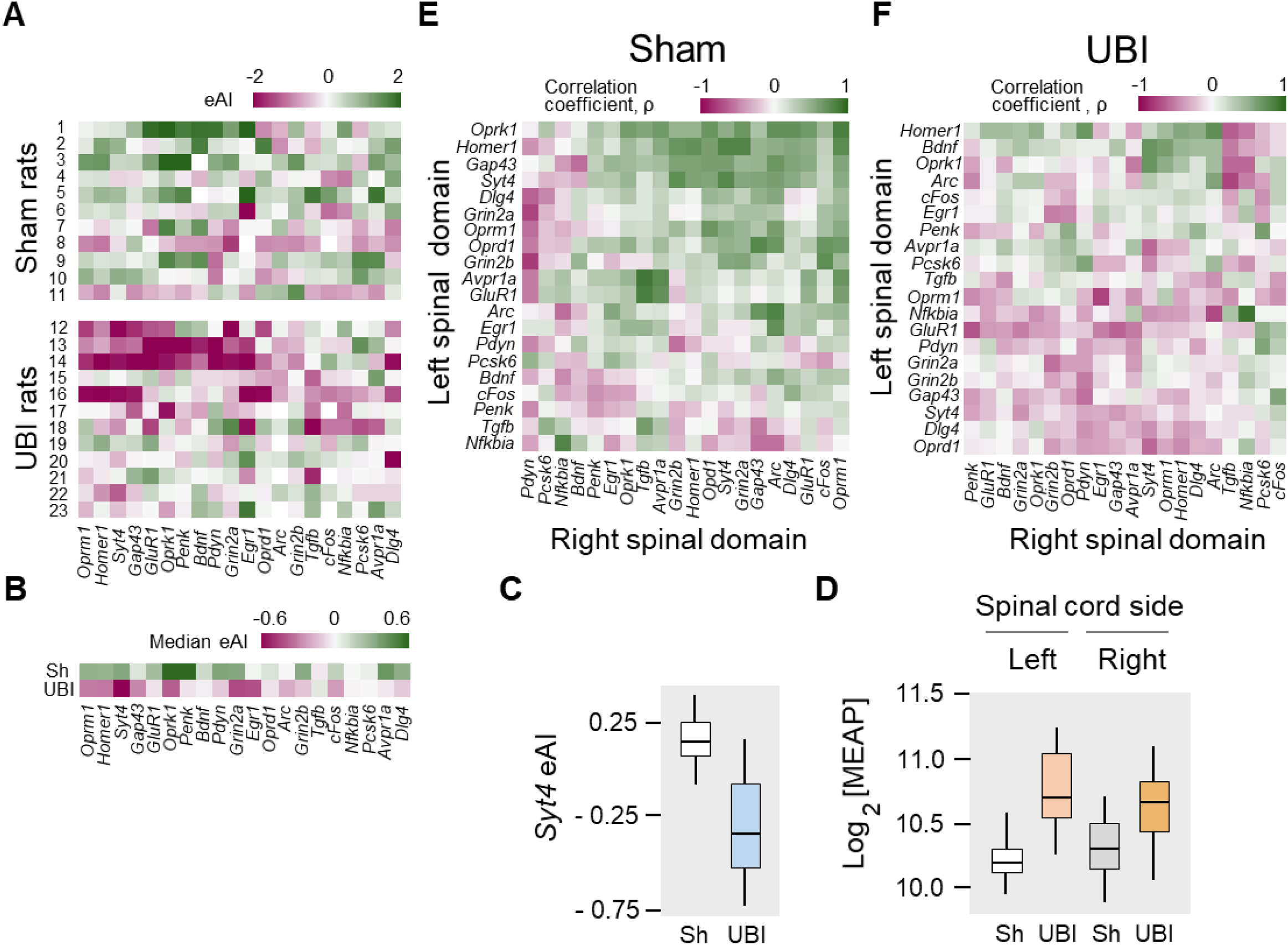
Expression of neuroplasticity and neuropeptide genes in the lumbar domains of rats exposed to the left UBI after complete spinal cord transection. The mRNA and peptide levels were analyzed in the ipsi- and contralesional halves of lumbar spinal cord isolated 3 h after the left UBI (n = 12) or left sham surgery (Sh; n = 11). (**A,B**) Heatmap for the expression asymmetry index (eAI=log_2_[Contra/Ipsi]) for each gene denoted for each rat individually, and as medians for rat groups. (**C**) UBI effects on the *Syt4* expression asymmetry index (median difference 0.38); Mann-Whitney test followed by Bonferroni correction: P_adjusted_ = 0.004. (**D**) UBI effects on the Met-enkephalin-Arg-Phe (MEAP) levels in the left (P_adjusted_ = 9×10^−4^; fold changes: 1.4) and right (P_un-djusted_ = 0.02; fold changes: 1.3) halves. Data is presented in fmole/mg tissue on the log_2_ scale as boxplots with medians. (**E,F**) Heatmap for Spearman correlation coefficients of expression levels between the left- and right lumbar halves for all gene pairs (inter-domain correlations) in rats with transected spinal cord that were exposed to sham surgery or UBI. **Source data 1.** The EXCEL source data file contains data for panels **A**-**C** of ***Figure 1***. **Source data 2.** The EXCEL source data file contains data for panel **D** of ***Figure 3***. **Figure supplement 1.** Genes analyzed and PCR Probes for their analysis. **Figure supplement 2.** Effects of the UBI on expression of neuroplasticity and neuropeptide genes, and the levels of opioid peptides in the lumbar spinal domains of the rats with transected spinal cord. **Figure supplement 3.** Heatmap for expression levels and intra-area correlations in the ipsi- and contralesional lumbar domains.

Gene co-expression patterns characterize regulatory interactions within and across tissues (Long et al., 2016). We examined whether left UBI performed after spinal transection induces changes in RNA–RNA intra-area correlations in the left and right halves of the lumbar spinal cord, and inter-area correlations between these halves. The proportion of intra-domain positive correlations, which dominated in rats with sham surgery, was reduced after the left UBI (Fisher’s Exact Test: all correlations in the right domain, P = 3×10^−5^; significant correlations in the left and right domains, P = 0.008 and 0.009, respectively (***Figure 3—figure supplement 3E-H***). The inter-domain gene-gene coordination strength was decreased after the left UBI (Wilcoxon signed-rank test; all and significant correlations: P = 4×10^−7^ and 3×10^−4^, respectively) (***Figure 3E,F***). Positive inter-domain correlations were predominant in rats with sham surgery (68%) in contrast to the UBI group (42%) (Fisher’s Exact Test: all and significant correlations, P = 6×10^−14^ and 0.004, respectively). We conclude that in rats with transected spinal cord the UBI robustly impairs coordination of expression of neuroplasticity and neuropeptide genes within and between the left and right halves of the lumbar spinal cord, and suggest that opioid neuropeptides may mediate spinal UBI effects. These experiments provide molecular evidence for the non-spinal cord mediated lateralized signaling from the injured brain to spinal neural circuits.

### UBI effects are mediated by neuroendocrine pathway

A non-spinal mechanism may operate through the neuroendocrine system by a release of pituitary hormones into the blood. Consistent with this hypothesis, no HL-PA was developed in hypophysectomized animals exposed to the left UBI after spinal transection (n = 8); the HL-PA median values and P_A_ were nearly identical to those in sham operated rats (n = 8) (***Figure 4A***; ***Figure 4—figure supplement 1A-E***). We next examined whether the left UBI stimulates the release of chemical factors, which may induce the development of HL-PA, into the blood. Serum that was collected 3 hours after performing a left UBI in rats with transected spinal cord was administered either centrally (into the cisterna magna; UBI serum, n = 13; sham serum, n = 7; ***Figure 4—figure supplement 1F-J***) or intravenously (UBI serum, n = 13; sham serum, n = 7; ***Figure 4B***; ***Figure 4—figure supplement 1K-O***) to rats after their spinalization. Serum administration by either route resulted in formation of HL-PA with its values and its probability similar to those induced by the UBI in rats with transected spinal cord. Remarkably, animals injected with serum from rats with left UBI displayed hindlimb flexion on the right side, which was the same as the flexion side in the donor rats (***Figure 4B—figure supplement 1F-O***). No HL-PA developed after administration of serum collected from rats with the left sham surgery. We conclude that the left UBI may stimulate a release of chemical factors from the pituitary gland, into the blood that induce HL-PA with contralesional flexion.

**Figure 4.**
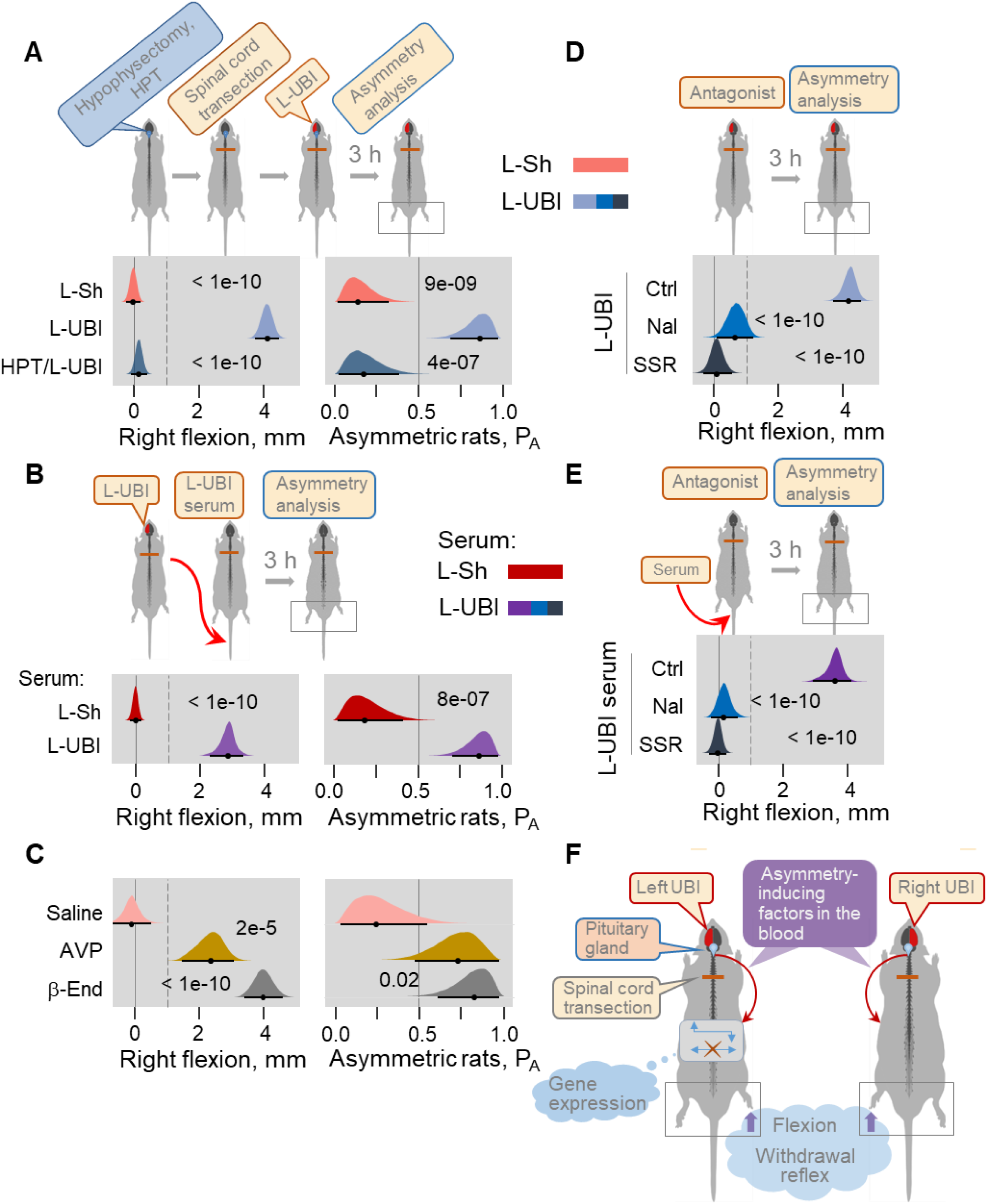
Neuroendocrine pathway mediating postural asymmetry formation in rats with transected spinal cord. (**A**) HL-PA in hypophysectomized (HPT; n = 8) and control (n = 12) rats with transected spinal cord 3 h after left UBI (L-UBI). Left sham surgery (L-Sh): n = 8. (**B**) HL-PA after intravenous administration of serum from rats with either left UBI (L-UBI serum) or left sham surgery (L-Sh serum) to rats with transected spinal cord (n = 13 and 7, respectively). (**C**) Induction of HL-PA by Arg-vasopressin (AVP) and β-endorphin (β-End) in rats with transected spinal cord. Synthetic β-endorphin or Arg-vasopressin (1 microgram and 10 nanogram / 0.3 ml saline / animal, respectively), or saline was administered intravenously to rats (n = 8, 7 and 4 rats), respectively after spinal cord transection. The HL-PA was analyzed in prone position 60 min after the injection under pentobarbital anesthesia. (**D**) Effect of naloxone (Nal, n = 6) or saline (n = 6), and SSR-149415 (SSR, n = 6) or vehicle (n = 5) on HL-PA 3 h after left UBI in rats with transected spinal cord. Vehicle and saline groups were combined into control group (Ctrl; n = 11). (**E**) Effect of naloxone (n = 6) or saline (n = 3) and SSR-149415 (n = 6) or vehicle (n = 3) on HL-PA 3 h after intravenous administration of the left UBI serum to rats with transected spinal cord. Ctrl; n = 6. In (**D,E**), naloxone (or saline) and SSR-149415 (or vehicle) were administered 0.5 and 3 h before HL-PA analysis, respectively. HL-PA values in millimeters (mm) and probability (P_A_) to develop HL-PA above 1 mm threshold (denoted by vertical dotted lines) are plotted as median, 95% HPDC intervals, and posterior distribution from Bayesian regression. Negative and positive HL-PA values are assigned to rats with the left and right hindlimb flexion, respectively. Significant asymmetry and differences between the groups: 95% HPDC intervals did not include zero value, and adjusted P-values were ≤ 0.05. Adjusted P is shown for significant differences identified by Bayesian regression. (**F**) Model for the humoral neuroendocrine side-specific signaling from the unilaterally injured brain to the lumbar spinal cord. In the rats with transected spinal cord, after the UBI the asymmetry inducing factors (neurohormones) may be released from the pituitary into the circulation, transported to their target sites and induce flexion of the contralesional hindlimb and asymmetric, contra vs. ipsilesional side specific changes in withdrawal reflexes and spinal gene expression patterns. **Source data 1.** The EXCEL source data file contains data for panels **A**-**E** of ***Figure 4***. **Figure supplement 1.** HL-PA formation in hypophysectomized rats exposed to the UBI and in control rats after administration of serum of the UBI animals. **Figure supplement 2.** Induction of HL-PA by Arg-vasopressin administered intracisternally to intact rats.

Multiple peptide factors inducing a side-specific hindlimb motor response were extracted from the brain, pituitary gland and serum of intact animals; several of them may be identical to endogenous peptides neurohormones (Bakalkin & Kobylyansky, 1989; Bakalkin et al., 1986; Chazov et al., 1981; Klement’ev et al., 1986). After central administration two factors, Arg-vasopressin and Leu-enkephalin induced HL-PA with right hindlimb flexion (for replication of Arg-vasopressin effects, see ***Figure 4—figure supplement 2***; peptide, n = 10; saline, n = 5/9). We here tested if β-endorphin and Arg-vasopressin, which both mostly are produced in and released into the circulation from the pituitary gland, may evoke asymmetric motor response after intravenous administration. Injection of these neurohormones but not saline to rats with transected spinal cord resulted in development of HL-PA with hindlimb flexion on the right side (***Figure 4C***; β-endorphin, n = 8; Arg-vasopressin, n = 7; saline, n = 4).

We next investigated whether opioid receptors and vasopressin receptor V1B, that is expressed in the pituitary gland (Roper et al., 2011), mediate formation of HL-PA in UBI rats or in animals treated with serum from UBI rats. Naloxone and SSR-149415, the opioid and vasopressin V1B receptor antagonists, respectively, administered to animals with transected spinal cord that were exposed to the left UBI (naloxone, n = 6; SSR-149415, n = 6; saline and vehicle, n = 11), or received serum from animals with left UBI (naloxone, n = 6; SSR-149415, n = 6; saline and vehicle, n = 6), inhibited HL-PA formation (***Figure 4D,E***). Thus activation of the receptors by the pituitary hormones β-endorphin and Arg-vasopressin released into the systemic circulation may be a necessary part of the hormone cascade mediating UBI effects on hindlimb motor circuits.

## Discussion

All three UBI effects in rats with transected spinal cord including development of HL-PA, asymmetry in nociceptive withdrawal reflexes, and asymmetric changes in gene expression patterns are mediated by a non-spinal endocrine pathway (***Figure 4F***). Encoding of information about the UBI and its laterality in a hormonal message, transmission of this message through the blood to its targets in peripheral endings of sensory neurons, dorsal root ganglia or non-neuronal (e. g. muscle or skin cells), and translation of this message into the left-right side specific response, are three stages of the phenomenon.

A bi-directional mechanism of the left-right side specific responses evoked by hormonal molecules circulating in the blood is a core of the non-spinal signaling pathway. This study in combination with our previous findings (Bakalkin & Kobylyansky, 1989; Bakalkin et al., 1986; Chazov et al., 1981) provide the principal evidence for such a mechanism. We demonstrated that peptide neurohormones and opioids administered intravenously, intrathecally or intracisternally induce HL-PA in rats with transected spinal cord. The striking finding was that the side of the flexed limb was dependent on the compound administered. Endogenous and synthetic κ-opioid agonists dynorphin and bremazocine, and endogenous mixed μ / δ-opioid agonist Met-enkephalin induced flexion of the left hindlimb (Bakalkin et al., 1989; Chazov et al., 1981). In contrast, β-endorphin, δ-agonist Leu-enkephalin and Arg-vasopressin caused the right limb to flex ((Bakalkin et al., 1989; Chazov et al., 1981; Klement’ev et al., 1986) and present study). Thus molecular signals circulating in blood were converted into the side-specific motor response suggesting that the opioid and vasopressin neurohormones may serve as chemical messengers transmitting information from an injured brain to peripheral tissues. The pituitary gland is the main source of opioid peptides and Arg-vasopressin in the bloodstream (Autelitano et al., 1989; Day & Akil, 1989). Naloxone and SSR-149415 blocked the UBI-induced formation of HL-PA demonstrating that the activation of the opioid receptors and the AVP V1b receptor is required for the signaling from the injured brain mediated through the pituitary gland to lumbar spinal circuits. A humoral brain-to-spinal cord signaling was earlier proposed in our study showing that administration of opioid peptide Met-enkephalin into the rostral portion of the spinal cord that was completely transected at the thoracic level, resulted in development of HL-PA with right hindlimb flexion (Bakalkin et al., 1986). Effects were likely mediated by circulated peptide hormones because serum collected from these rats and injected to intact animals also induced hindlimb flexion on the right side.

Hypothetically, in animals with the T2-3 transected spinal cord asymmetric effects of the UBI on hindlimb motor functions may be mediated by the sympathetic outflow from the upper thoracic segments to hindlimb muscle vasculature. However, projections of the preganglionic neurons located above the T5 level pass superiorly along the paravertebral sympathetic trunks, those located below this level pass inferiorly, and the preganglionic fibers for lower limbs are derived from the lower three thoracic and upper lumbar spinal segments, i.e., 8 segments below the transection level. Furthermore, the sympathetic nervous system has a limited capacity to independently regulate blood flow to the left and right hindlimbs (Lee et al., 2007). In the absence of published experimental support for a putative sympathetic mechanism these observations suggest that the UBI-induced asymmetric sympathetic effects on hindlimb vasculature may be limited, if any. Our findings do not rule out this possibility, but provide unequivocal evidence for the endocrine signaling.

Lesion of the hindlimb sensorimotor area may also affect forelimbs that in turn may alter the hindlimb burden. While these effects may contribute to hindlimb motor performance in intact animals, e.g. in locomotor assays, they are apparently negligible, if any, in our experimental design. Lesion of this area did not produce noticeable forelimb postural asymmetry before and after complete spinal transection (Zhang et al., 2020) while spinal cord transection performed before the UBI eliminated neural interactions between fore and hindlimbs.

The side-specific effects suggest that spinal neural circuits regulating the left and right hindlimb muscles differ in sensitivity towards opioid and Arg-vasopressin neurohormones (Bakalkin et al., 1989; Chazov et al., 1981; Klement’ev et al., 1986). Lateralization of receptors for these peptides in the spinal cord or peripheral tissues, along with anatomical asymmetry of spinal sensorimotor circuits may be a basis for different regulation of the left- or right-side functional responses. The asymmetric organization of the spinal cord was demonstrated in physiological, anatomical and molecular studies (de Kovel et al., 2017; Hultborn & Malmsten, 1983; Kononenko et al., 2017; Nathan et al., 1990; Ocklenburg et al., 2017; Zhang et al., 2020). Activity of mono- and polysynaptic segmental reflexes is higher on the right-compared to left-side in rats and cats (Hultborn & Malmsten, 1983; Zhang et al., 2020). Three-quarters of spinal cords are asymmetric with larger right side (Nathan et al., 1990). The genes for the opioid receptors and the opioid peptides are asymmetrically expressed in the cervical spinal cord (Kononenko et al., 2017). All three opioid receptors are lateralized to the left while in different proportions. Their expression was coordinated between the dorsal and ventral domains but with different patterns on the left and right spinal sides.

In conclusion, this study describes a novel phenomenon, the side-specific endocrine mechanism that mediates asymmetric effects of unilateral brain injury on hindlimb motor deficits (***Figure 4F***). The humoral pathway and the descending neural tracts may represent complementary routes for signaling from the brain to the spinal cord. Analysis of features and proportion of sensorimotor deficits transmitted by neurohormonal signals versus those mediated by neural pathways in animal models and patients after stroke and traumatic brain injury should facilitate new therapeutic discoveries. From a biological standpoint, the mechanism may serve to maintain a balance between the left–right processes in bilaterally symmetric animals.

## Figure supplements

**Figure 1—figure supplement 1.**
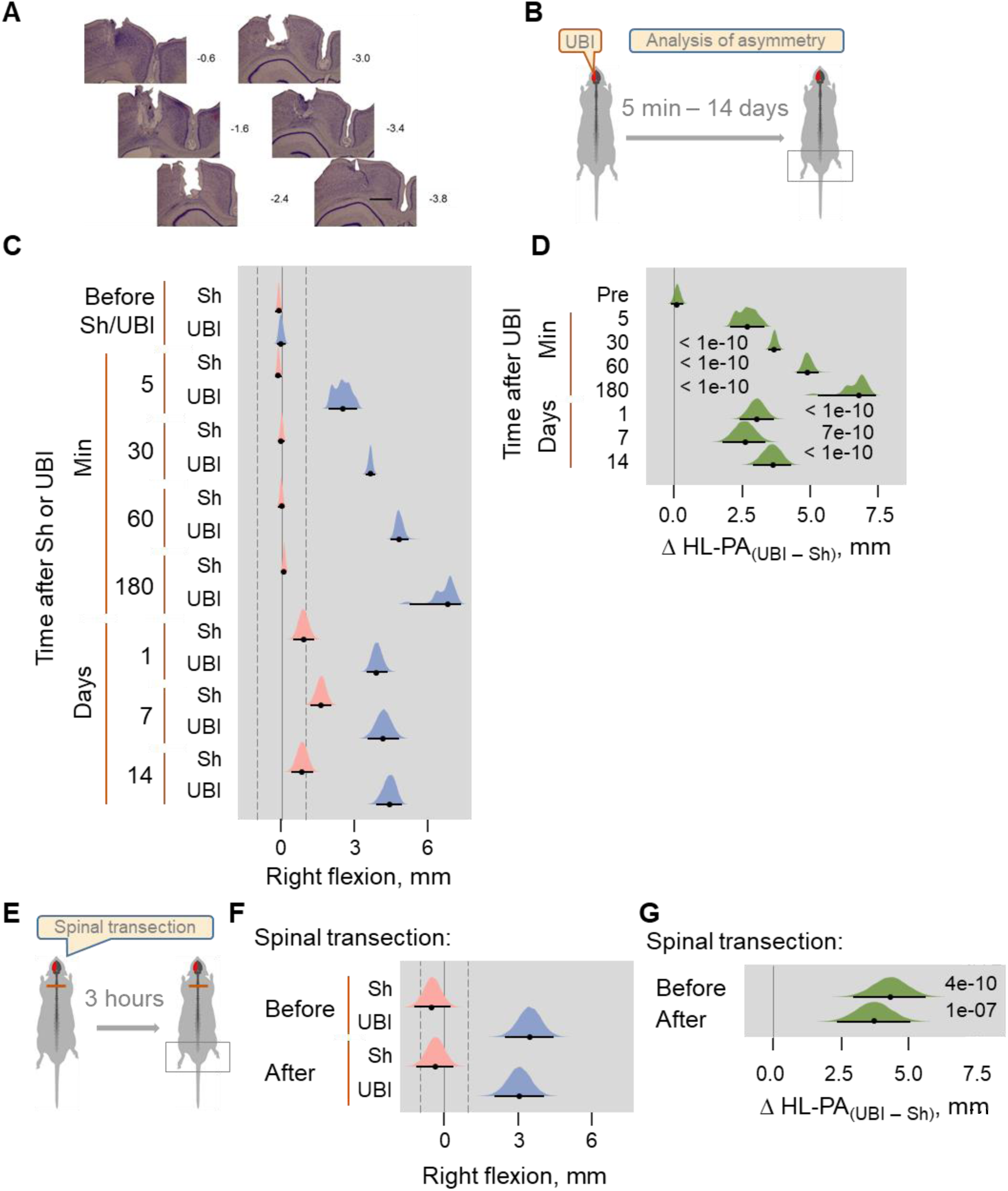
UBI-induced HL-PA formation and its fixation after complete spinal cord transection at the T2-3 level. (**A**) Histological verification of the size and location of UBI resulting from a stereotaxic aspiration of brain tissue at the following coordinates: 0.5 – 4.0 mm posterior to the bregma and 1.8 – 3.8 lateral to the midline. Series of Nissl-stained coronal brain sections show representative lesion of the rat sensorimotor cortex. The numbers on the right indicate distances (in mm) from bregma. Scale bar = 1 mm. Sections like these were obtained from all rats used in this study and the general distribution pattern and the extent of the lesions observed were reliably similar across subjects. (**B,E**) Experimental designs. (**C,D**) The HL-PA was analyzed in the prone position before and after left UBI (UBI) or left sham surgery (Sh) within 5 min and at the 30, 60 and 180 min time points under pentobarbital anesthesia (UBI, n = 8, Sh, n = 7); and at the 1, 7 and 14-day time points under isoflurane anesthesia (UBI, n = 12; Sh, n = 8). (**E-G**) In the rat subgroup on the 4^th^ day after left UBI (n = 5) or left sham surgery (n = 10), the spinal cord was transected under pentobarbital anesthesia and HL-PA was analyzed 3 h later. (**C,F**) Effects of UBI on the formation of HL-PA (expressed in millimeters, mm). Negative and positive HL-PA values are assigned to rats with the left and right hindlimb flexion, respectively. (**D,G**) Differences in HL-PA [Δ HL-PA_(UBI – Sh)_ in millimetrers] between UBI and sham goups. Medians, 95% HPDC intervals and densities from Bayesian sampler are plotted. Significant contrasts between the groups: 95% HPDC intervals did not include zero value, and adjusted P-values were ≤ 0.05. Adjusted P is shown for significant differences identified by Bayesian regression.

**Figure 1—figure supplement 2.**
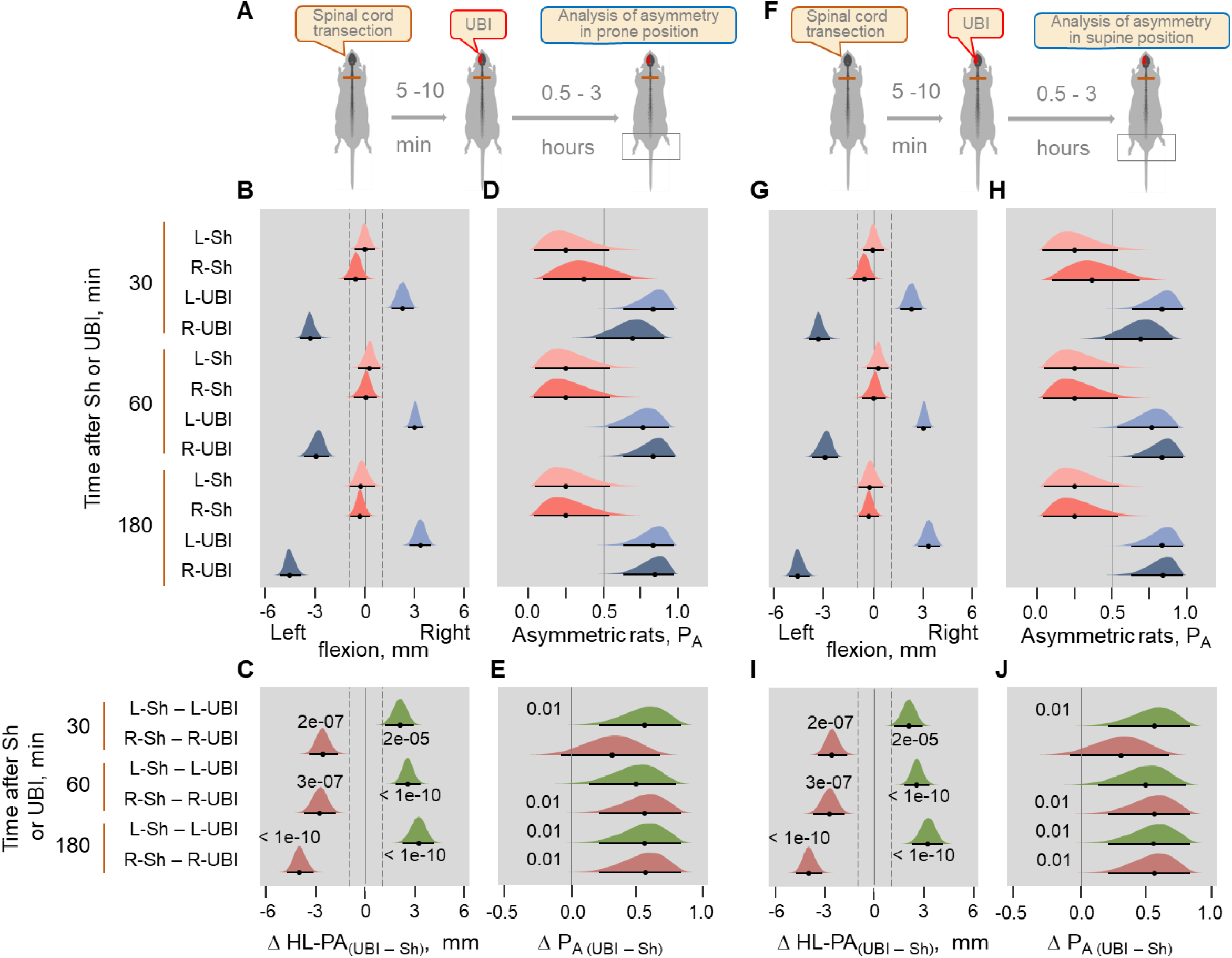
Time-course of HL-PA formation analyzed in prone (A-E) and supine (F-J) positions after the left-side (L-UBI) and right-side (R-UBI) UBI in Wistar rats with transected spinal cord. (**A,F**) Experimental designs. The UBI was conducted 5 -10 min after complete spinal cord transection at the T2-T3 level. The HL-PA was analyzed under pentobarbital anesthesia 30, 60 and 180 min after the brain injury. Data presented for the 180 min time point on ***Figure 1F*** are shown for comparison. The left UBI and right UBI groups consisted of 9 rats each, while the control groups consisted of 4 left- (L-Sh) and 4 right- (R-Sh) sham animals. (**B,D,G,H**) Effects of UBI on the formation of HL-PA (expressed in millimeters, mm), and on the probability to develop HL-PA (P_A_). The rats with the HL-PA magnitude above the 1 mm threshold (that is 99^th^ HL-PA magnitude percentile in sham groups; shown by vertical dashed lines) were defined as asymmetric. Negative and positive HL-PA values are assigned to rats with the left and right hindlimb flexion, respectively. (**C,E,I,J**) Differences in the HL-PA [Δ HL-PA_(UBI – Sh)_ in millimetrers] and in the the probability to develop HL-PA [Δ P_A (UBI – Sh)_] between UBI and sham surgery groups. Medians, 95% HPDC intervals and densities from Bayesian sampler are plotted. Significant contrasts between the groups: 95% HPDC intervals did not include zero value, and adjusted P-values were ≤ 0.05. Adjusted P is shown for significant differences identified by Bayesian regression.

**Figure 1—figure supplement 3.**
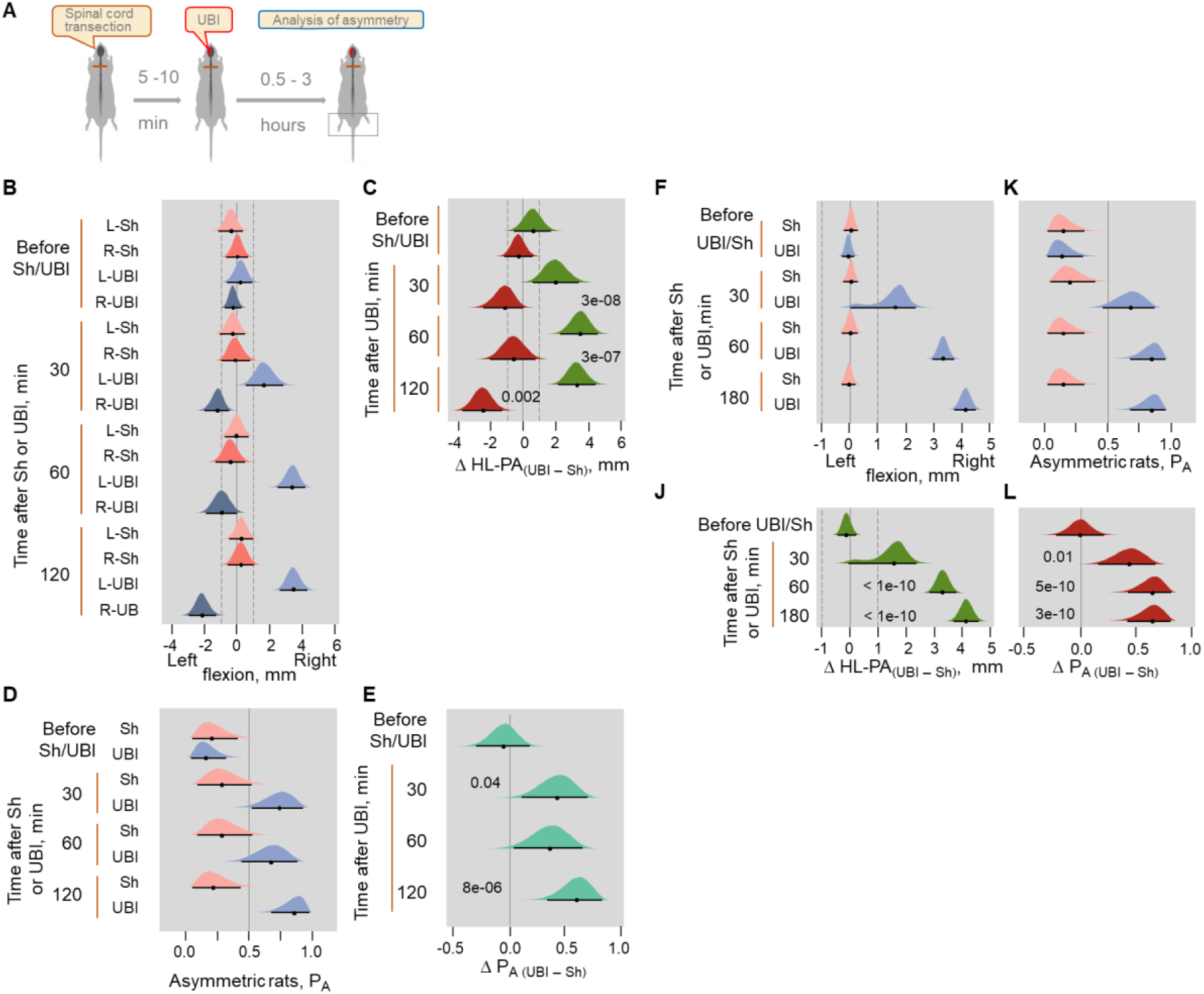
Replication experiments 1 (B-E) and 2 (F-L). (**A**) Experimental design. Induction of HL-PA by the UBI in rats with transected spinal cord. (**B**-**E**) Experiment 1; the HL-PA in Sprague Dawley rats was analyzed in prone position under pentobarbital anesthesia 30, 60 and 120 min after the left (n = 5) or right (n = 6) UBI, and the left (n = 5) or right (n = 5) sham surgery performed after complete spinal cord transection at the T2-T3 level. (**D,E**) The left and right UBI groups, as well as the left and right sham surgery groups were combined into the UBA and sham groups, respectively. (**F-L**) Experiment 2; the HL-PA in Wistar rats was analyzed in prone position under pentobarbital anesthesia 30, 60 and 180 min after the left UBI (n = 12) and the left sham surgery (n = 11) performed after complete spinal cord transection at the T2-T3 level. Negative and positive HL-PA values are assigned to rats with the left and right hindlimb flexion, respectively. Differences in the HL-PA [Δ HL-PA _(UBI – Sh)_ in millimetrers] and in the probability to develop HL-PA [Δ P_A (UBI – Sh)_] between UBI and sham surgery groups. Medians, 95% HPDC intervals and densities from Bayesian sampler are plotted. Significant contrasts between the groups: 95% HPDC intervals did not include zero value, and adjusted P-values were ≤ 0.05. Adjusted P is shown for significant differences identified by Bayesian regression.

**Figure 2—figure supplement 1.**
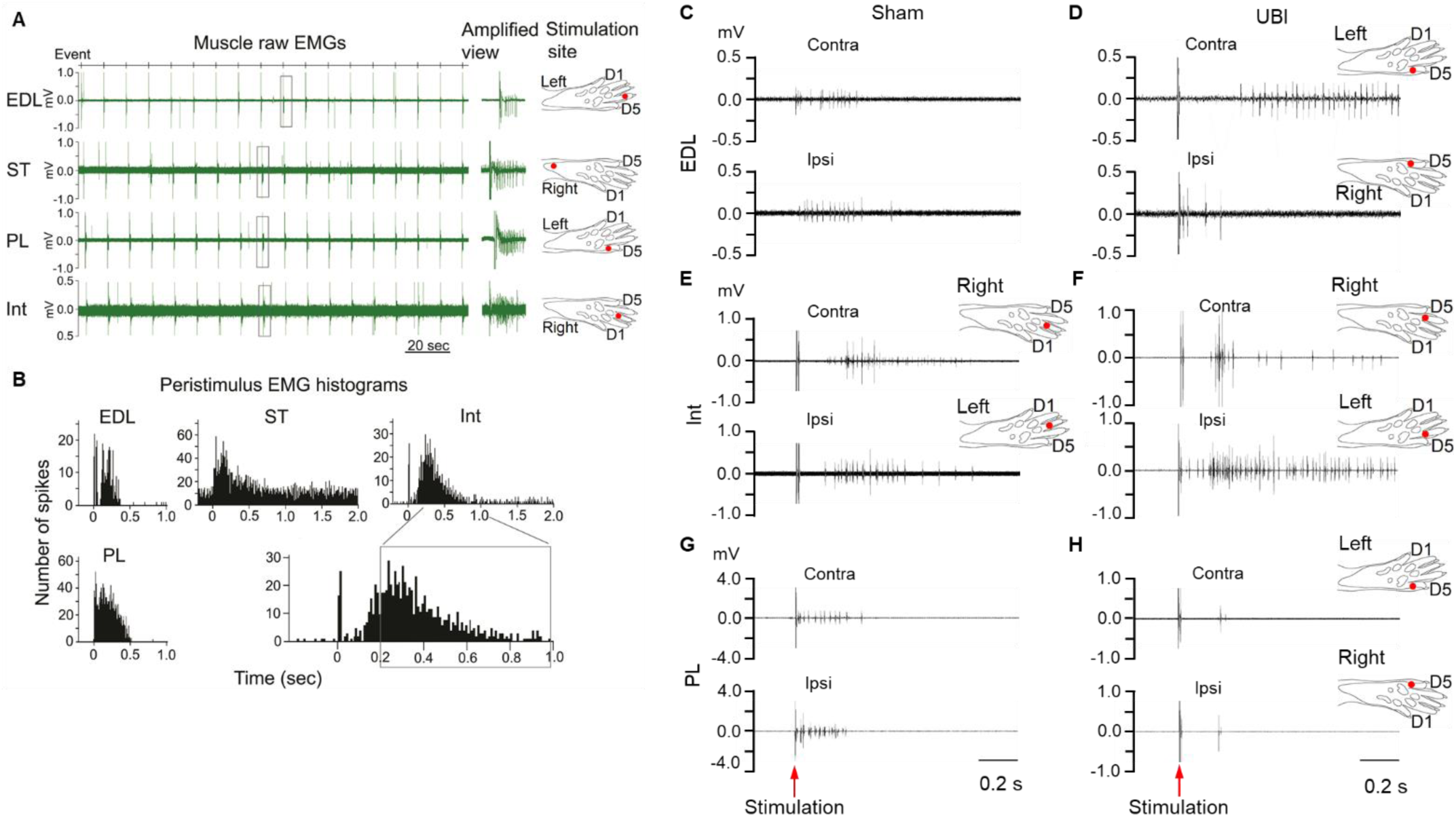
Effects of the UBI on hindlimb nociceptive withdrawal reflexes. (**A,B**) Representative EMG examples and respective stimulation sites of hindlimb muscles of the anesthetized rats with transected spinal cord that were exposed to the UBI. (**A**) Left panel: EMG responses of the extensor digitorum longus (EDL), semitendinosus (ST), peroneus longus (PL) and the forth interosseous (Int) muscles to 18 electrical stimulations. Middle panel: Amplified view of EMG spikes from the regions delimited by the rectangles on the left panel. Right panel: Stimulation site for each muscle. (**B**) Peristimulus histogram of these muscles from 16 stimulations (from 2^nd^ to 17^th^). The area delimited by the rectangle shows the time window (0.2 – 1.0 s) that was analysed statistically. (**C-H**) Representative examples of EMG responses of extensor digitorum longus (**C,D**), interosseous (**E,F**) and peroneus longus (**G,H**) muscles of the contra- and ipsilesional hindlimbs in rats with transected spinal cord that were exposed to UBI or sham surgery. Stimulation of the left and right hindlimbs of the UBI rat with the same current parameters induced larger EMG responses of the extensor digitorum longus muscle on the contra-compared to the ipsilesional side (**D**), and the interosseous muscle on the ipsi-compared to the contralesional side (**F**). Each of three muscles of sham rats (**C,E,G**) and of peroneus longus muscle of the UBI rat (**H**) demonstrated similar responses on both sides.

**Figure 2—figure supplement 2.**
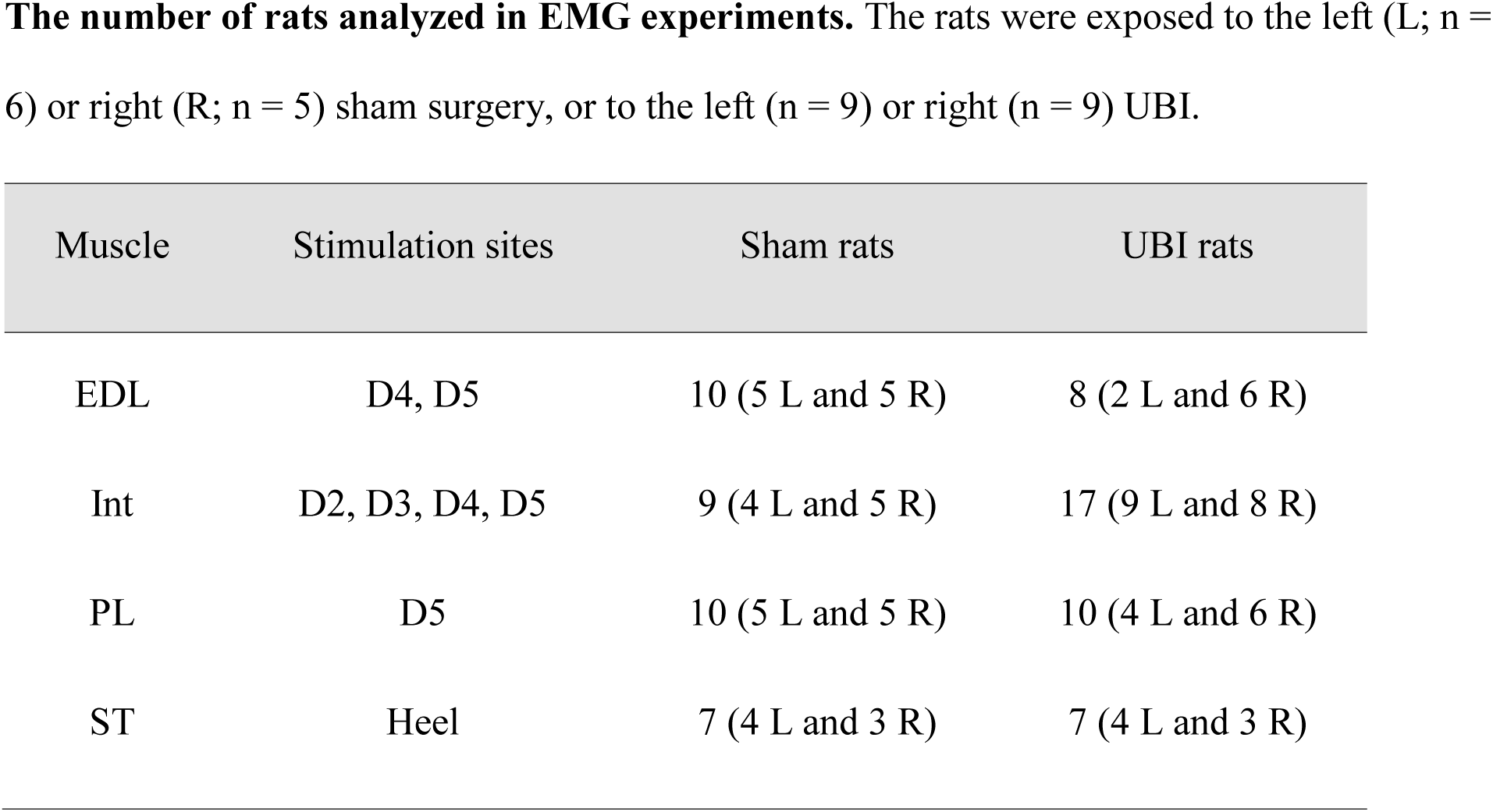
The number of rats analyzed in EMG experiments. The rats were exposed to the left (L; n = 6) or right (R; n = 5) sham surgery, or to the left (n = 9) or right (n = 9) UBI.

**Figure 3—figure supplement 1.**
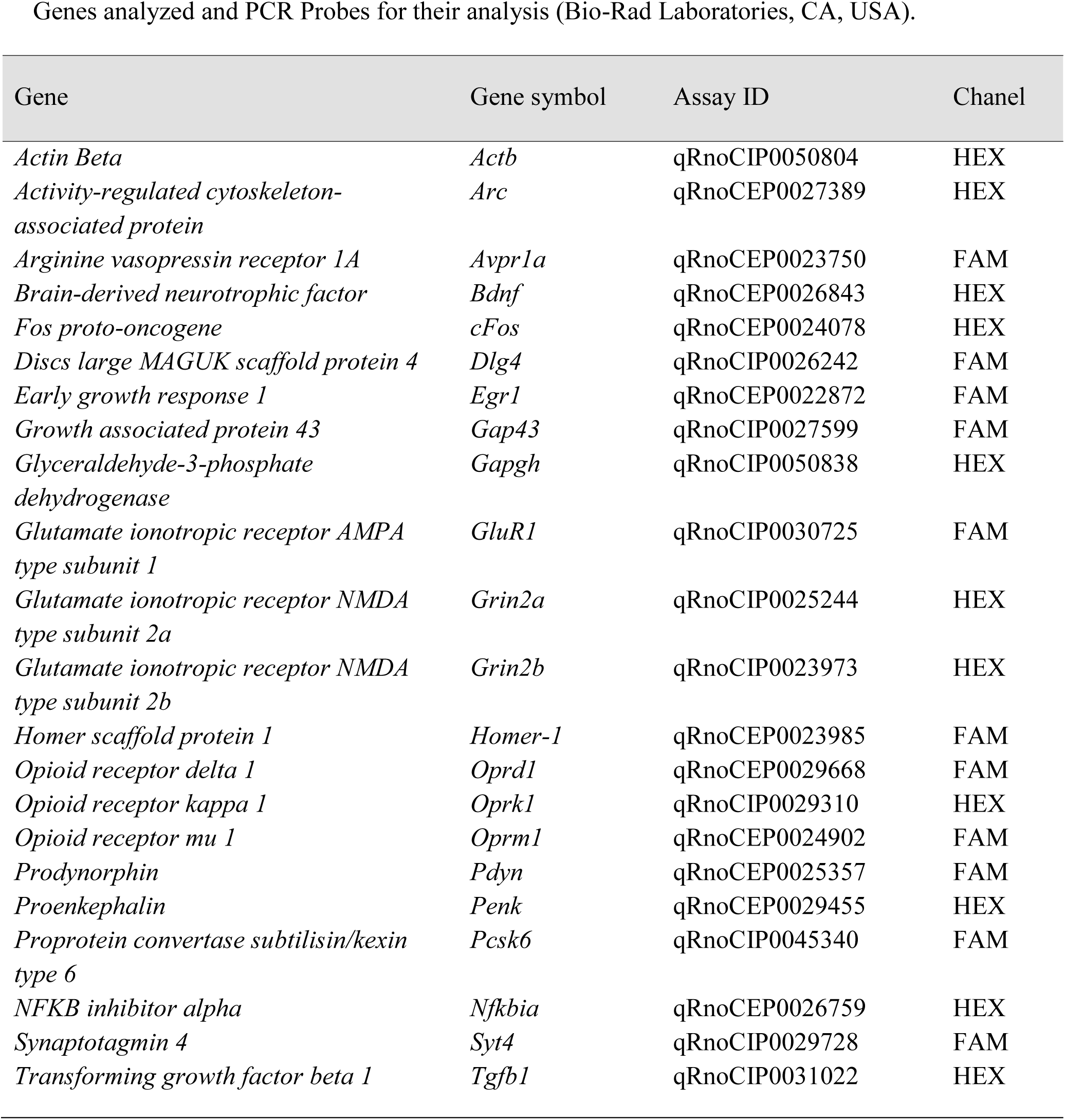
Genes analyzed and PCR Probes for their analysis (Bio-Rad Laboratories, CA, USA).

**Figure 3—figure supplement 2.**
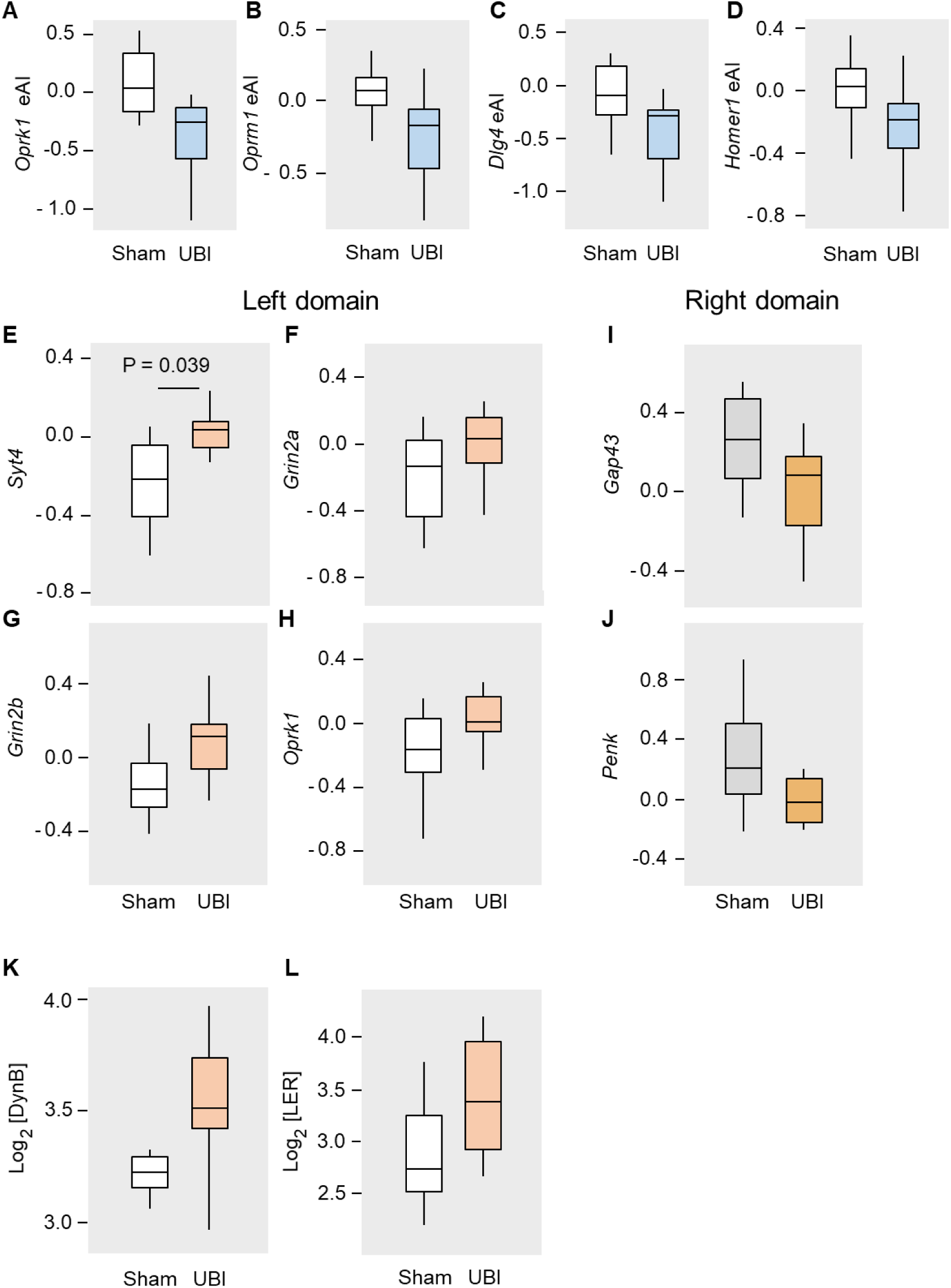
Effects of the UBI on expression of neuroplasticity and neuropeptide genes (A-J), and the levels of opioid peptides (K,L) in the lumbar spinal domains of the rats with transected spinal cord. The rats with spinal cord transection at the T2-T3 level were exposed the left UBI (n = 12) or left sham surgery (n=11), and the mRNA levels were analyzed in the ipsi- (left) and contralesional (right) domains of lumbar spinal cord isolated 3 h after the injury. (**A-D**) The median expression asymmetry index (eAI = log_2_[Contra/Ipsi], where Contra and Ipsi were the levels in the contra- and ipsilesional lumbar domains) for each of 20 genes was compared individually between the UBI and sham groups using Mann-Whitney test. Data is presented as boxplots with medians. The expression asymmetry index for the *Oprk1* (median difference 0.29; P_un-adjusted_ = 0.009), *Oprm1* (median difference 0.24; P _un-adjusted_ = 0.032), *Dlg4* (median difference 0.18; P _un-adjusted_ = 0.032) and *Homer1* (median difference 0.19; P _un-adjusted_ = 0.037) are shown. (**E-J**) Significance of the UBI-induced changes in expression of *Syt4* (1.19-fold; P_adjusted_ = 0.039), *Grin2a* (1.18-fold; P _un-adjusted_ = 0.04), *Grin2b* (1.22-fold; P _un-adjusted_ = 0.01) and *Oprk1* (1.13-fold; P _un-adjusted_ = 0.03) in the left lumbar domain; and *Gap43* (1.13-fold; P _un- adjusted_ = 0.04) and *Penk* (1.17-fold; P _un-adjusted_ = 0.03) in the right lumbar domain of the rats with transected spinal cord is shown. The mRNA levels of 20 genes were compared separately for the left and right halves of the lumbar spinal cord between the rats exposed to UBI and sham surgery using Mann-Whitney test followed by Bonferroni correction for a number of tests (n = 40). Data is presented on the log_2_ scale as boxplots with medians. (**K,L**) Effects of the UBI on the levels of opioid peptides Dynorphin B (DynB) (1.2-fold; Mann-Whitney test: unadjusted P = 0.03) and Leu-enkephalin-Arg (LER) (1.6-fold; Mann-Whitney P _un-adjusted_ = 0.02) in the ipsilesional (left) domain. Data is presented in fmol/mg tissue on the log_2_ scale as boxplots with medians.

**Figure 3—figure supplement 3.**
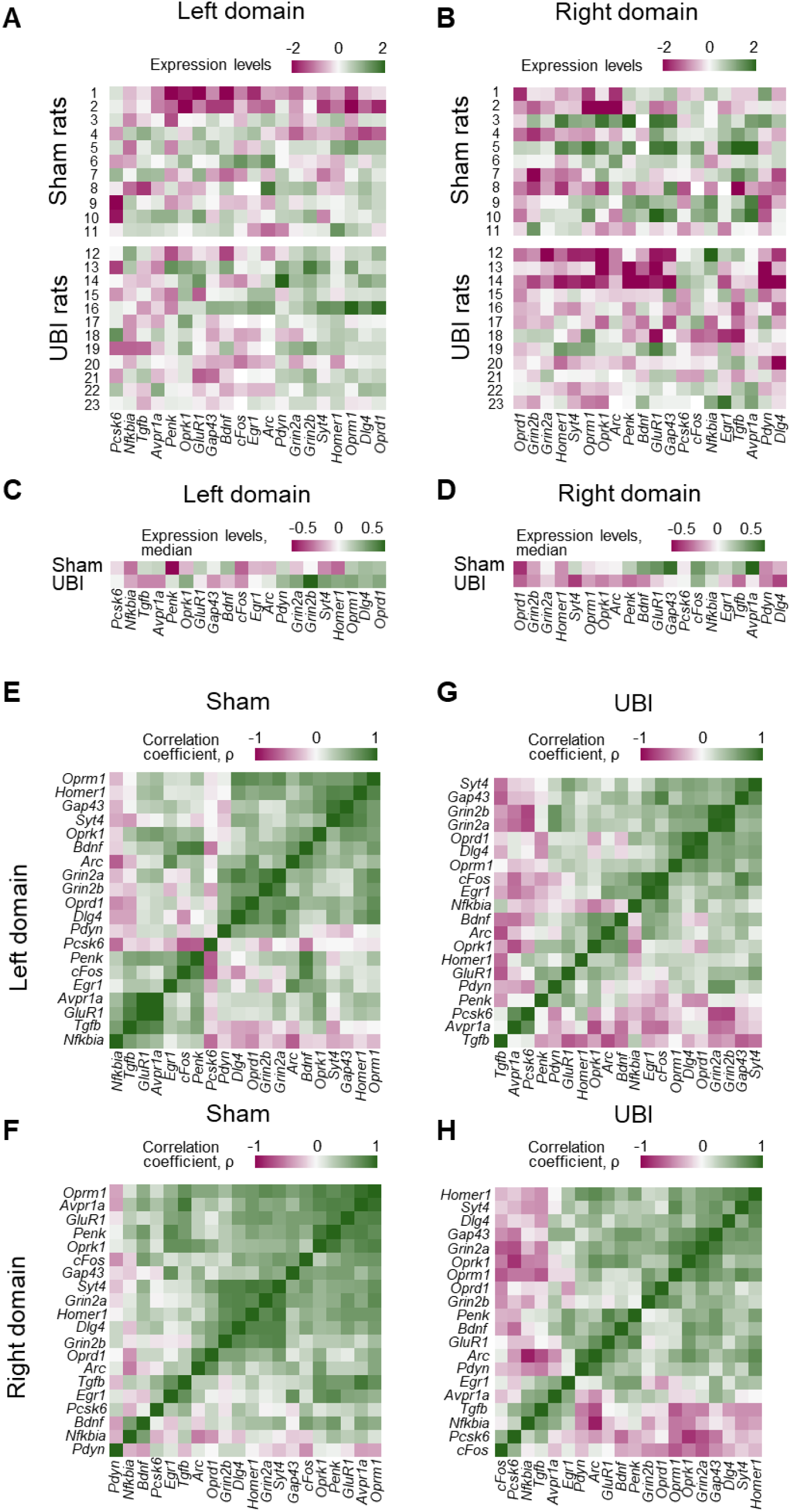
Heatmap for expression levels (A-D) and intra-area correlations (E-H) in the left (ipsilesional) (A,C,E,G) and right (contralesional) (B,D,F,H) lumbar domains. (**A-D**) Expression levels for each gene are denoted as (0,1)-standardized values for each sham and UBI rat individually (**A,B**), and as medians for the sham and UBI groups (**C,D**). The data is presented on the log_2_ scale. In the left domain 15 out of 20 analyzed genes have higher median expression in the UBI group (sign test: P = 0.041), while in the right domain for 17 out of 20 genes median expression is higher in the sham group (sign test: P = 0.003). (**E,H**) Heatmap for (0,1)-standardized coefficients of Spearman correlations in expression levels for all gene pairs in the rats exposed to left sham surgery (**E,F**) or left UBI (**G,H**). Differences between the UBI and sham groups in the proportion of positive and negative intra-area correlations were analyzed by the Fisher’s Exact Test: for all correlations in the right domain: P = 2.7×10^−5^; for significant correlations in the left and right domains, P = 8.1×10^−3^ and 8.9×10^−3^, respectively.

**Figure 4—figure supplement 1.**
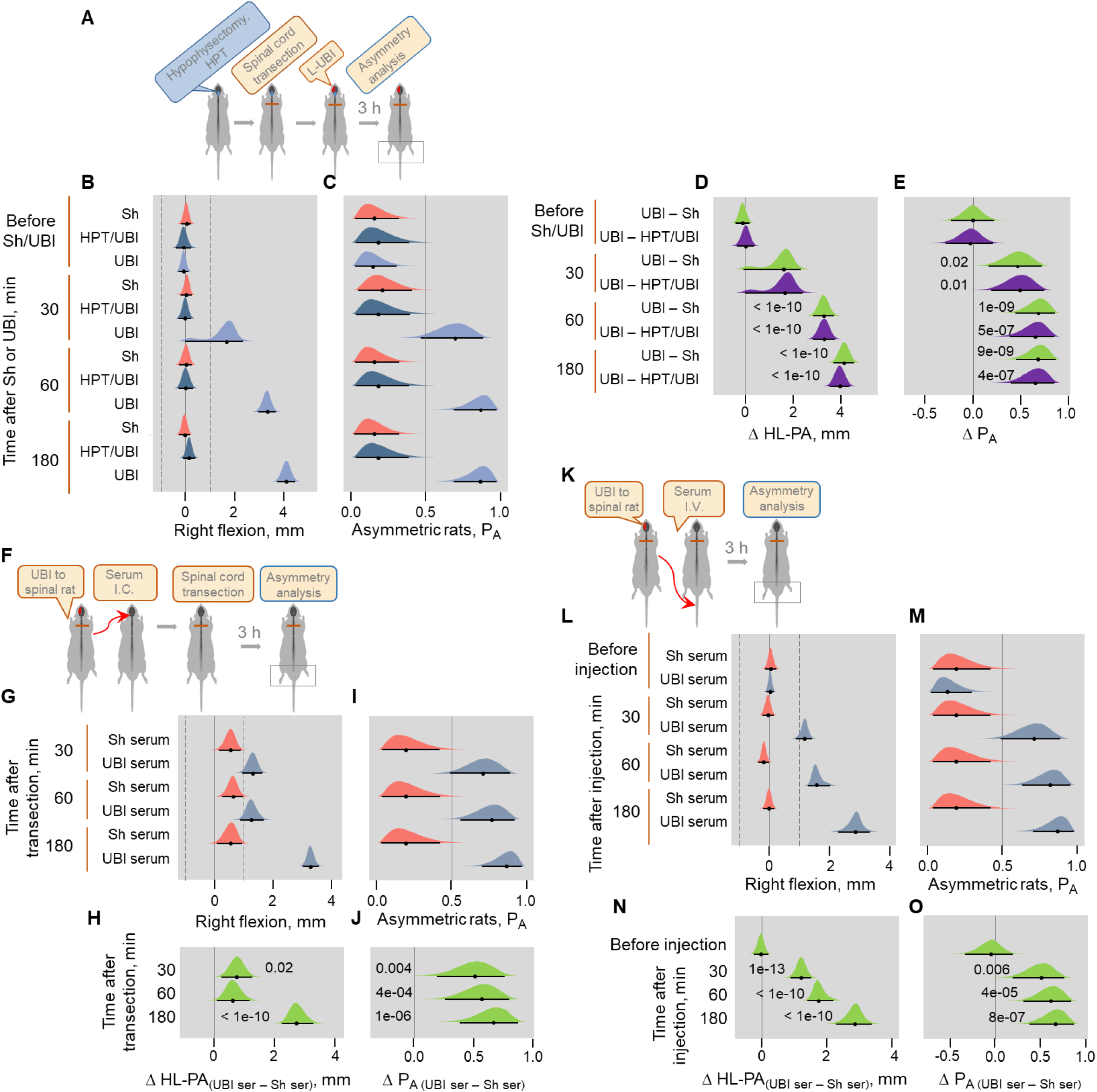
HL-PA formation in hypophysectomized rats exposed to the UBI, and in rats after administration of serum of the UBI animals. (**A-E**) The left UBI in hypophysectomized (n = 8) and control “intact” (n = 12) rats, and left sham surgery in control “intact” rats (n = 8) were performed after complete spinal cord transection at the T2-T3 level. (**F-J**) Induction of HL-PA by serum of the UBI animals. Serum collected from the rats with transected spinal cord 3 hours after their exposure to the left UBI (UBI serum) or sham surgery (Sh serum) was administered to the cisterna magna (I.C.; 5 microliters / rat) to intact rats. The spinal cord was transected at the T2-T3 level 10-15 min after injection of the UBI (n = 13) or sham (n = 7) serum, and the HL-PA was analyzed in prone position 30, 60 and 180 min after the transection under pentobarbital anesthesia. (**K-O**) Time-course of HL-PA formation after intravenous administration of the left UBI serum to rats with transected spinal cord. HL-PA was analyzed after administration I.V. of serum collected from the rats exposed to the left-side UBI (UBI serum; n = 13) or sham surgery (Sham serum; n = 7), to the rats after their spinalization at the T2-T3 level. Data for the 180 min time point presented on ***Figure 4B*** are shown for comparison. Negative and positive HL-PA values are assigned to rats with the left and right hindlimb flexion, respectively. Differences in the HL-PA [Δ HL-PA _(UBI – Sh)_ in millimetrers] and in the probability to develop HL-PA [Δ P_A (UBI – Sh)_] between UBI and sham surgery groups. Medians, 95% HPDC intervals and densities from Bayesian sampler are plotted. Significant contrasts between the groups: 95% HPDC intervals did not include zero value, and adjusted P-values were ≤ 0.05. Adjusted P is shown for significant differences identified by Bayesian regression.

**Figure 4—figure supplement 2.**
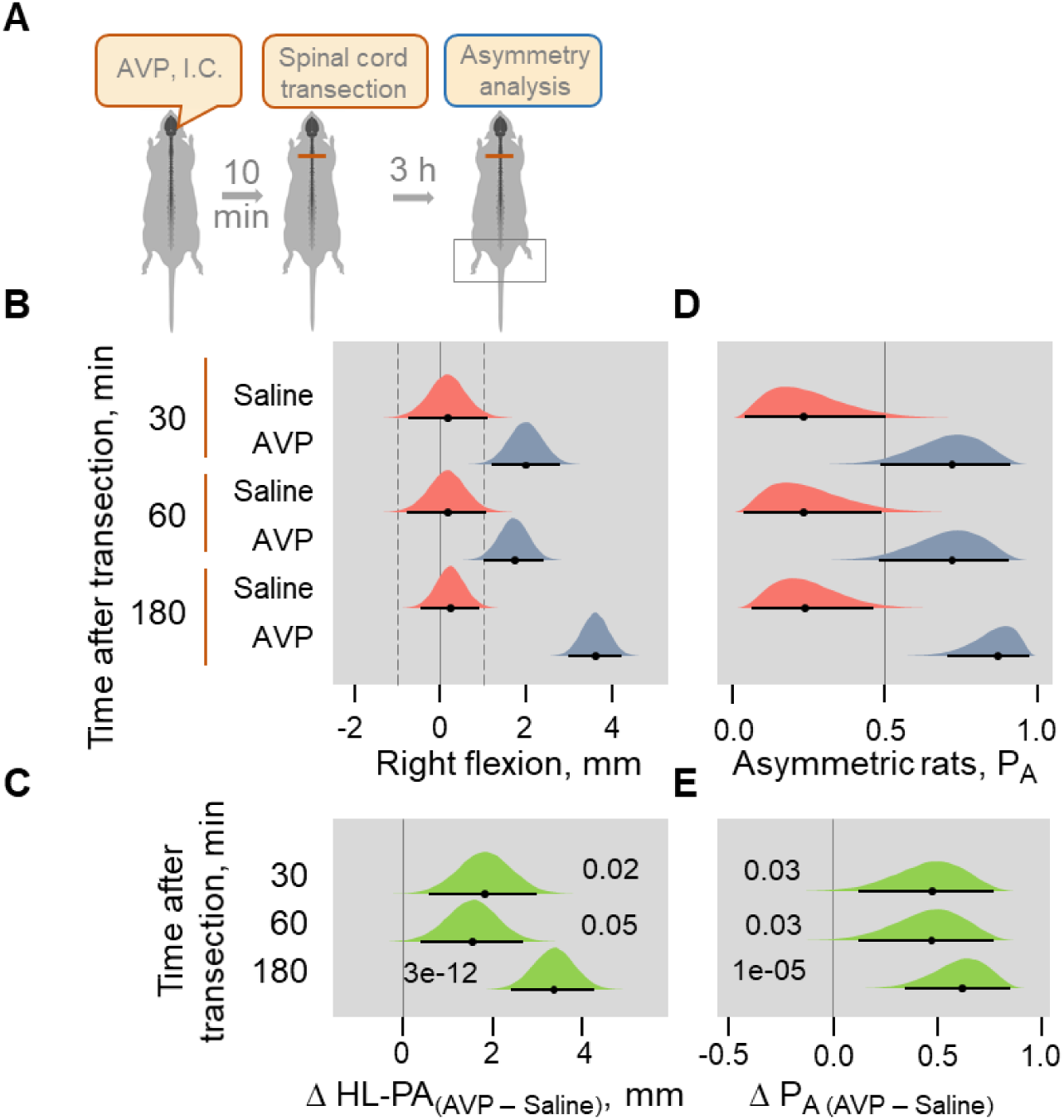
Induction of HL-PA by Arg-vasopressin (AVP) administered intracisternally to rats with transected spinal cord. HL-PA formation was analyzed in prone position under pentobarbital anesthesia 30, 60 and 180 min after AVP (10 nanograms in 5 microliters / rat; n = 10 at the 30 and 60 min time points, and n = 22 at the 180 min time point) or saline (n = 5 at the 30 and 60 min time points, and n = 9 rats at the 190 min time point) administration followed in 10-15 min by spinal cord transection. Negative and positive HL-PA values are assigned to rats with the left and right hindlimb flexion, respectively. Differences in the HL-PA [Δ HL-PA _(UBI – Sh)_ in millimetrers] and in the probability to develop HL-PA [Δ P_A (UBI – Sh)_] between UBI and sham surgery groups. Medians, 95% HPDC intervals and densities from Bayesian sampler are plotted. Significant contrasts between the groups: 95% HPDC intervals did not include zero value, and adjusted P-values were ≤ 0.05. Adjusted P is shown for significant differences identified by Bayesian regression.

## Materials and methods

### Key resources table

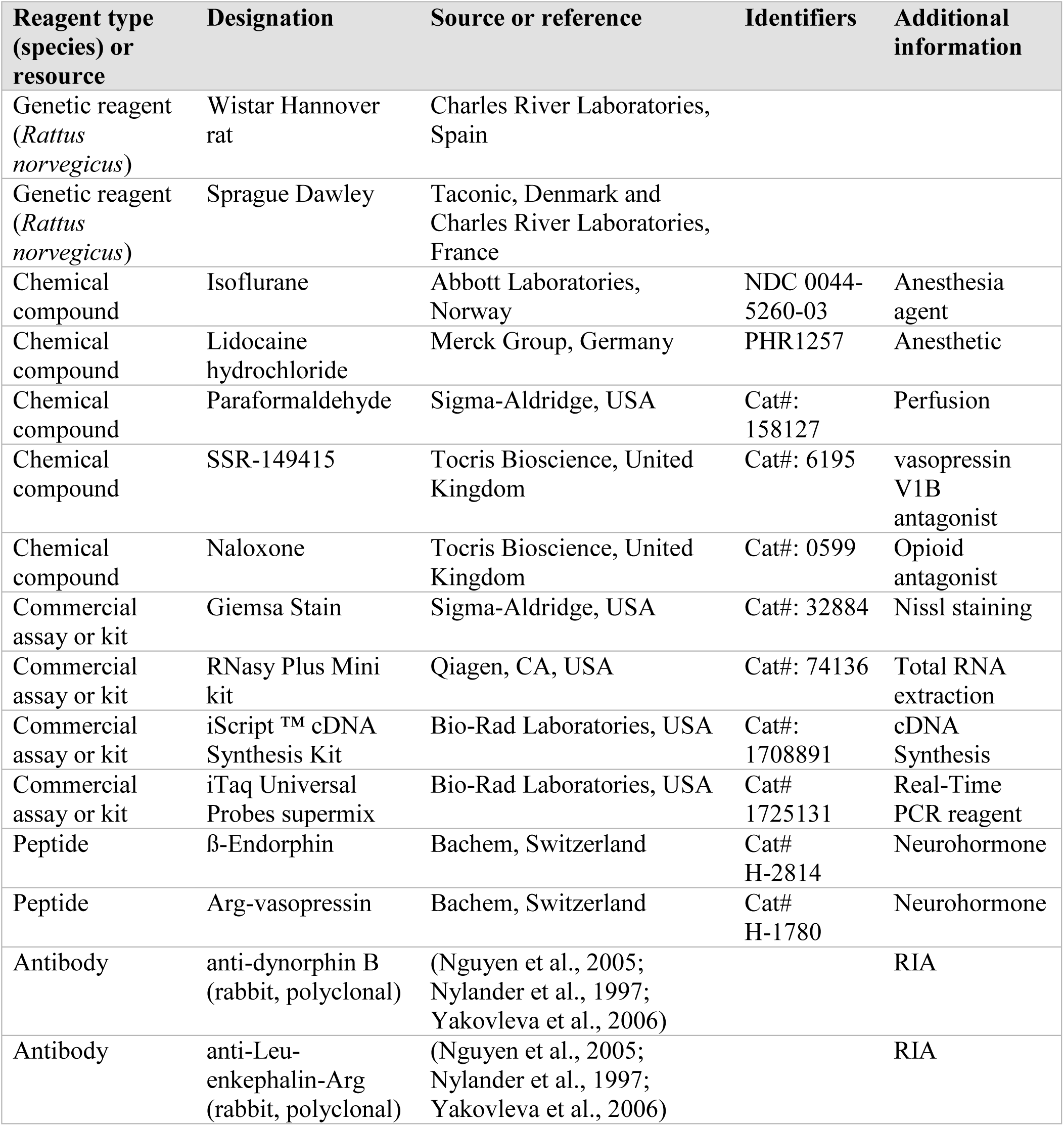
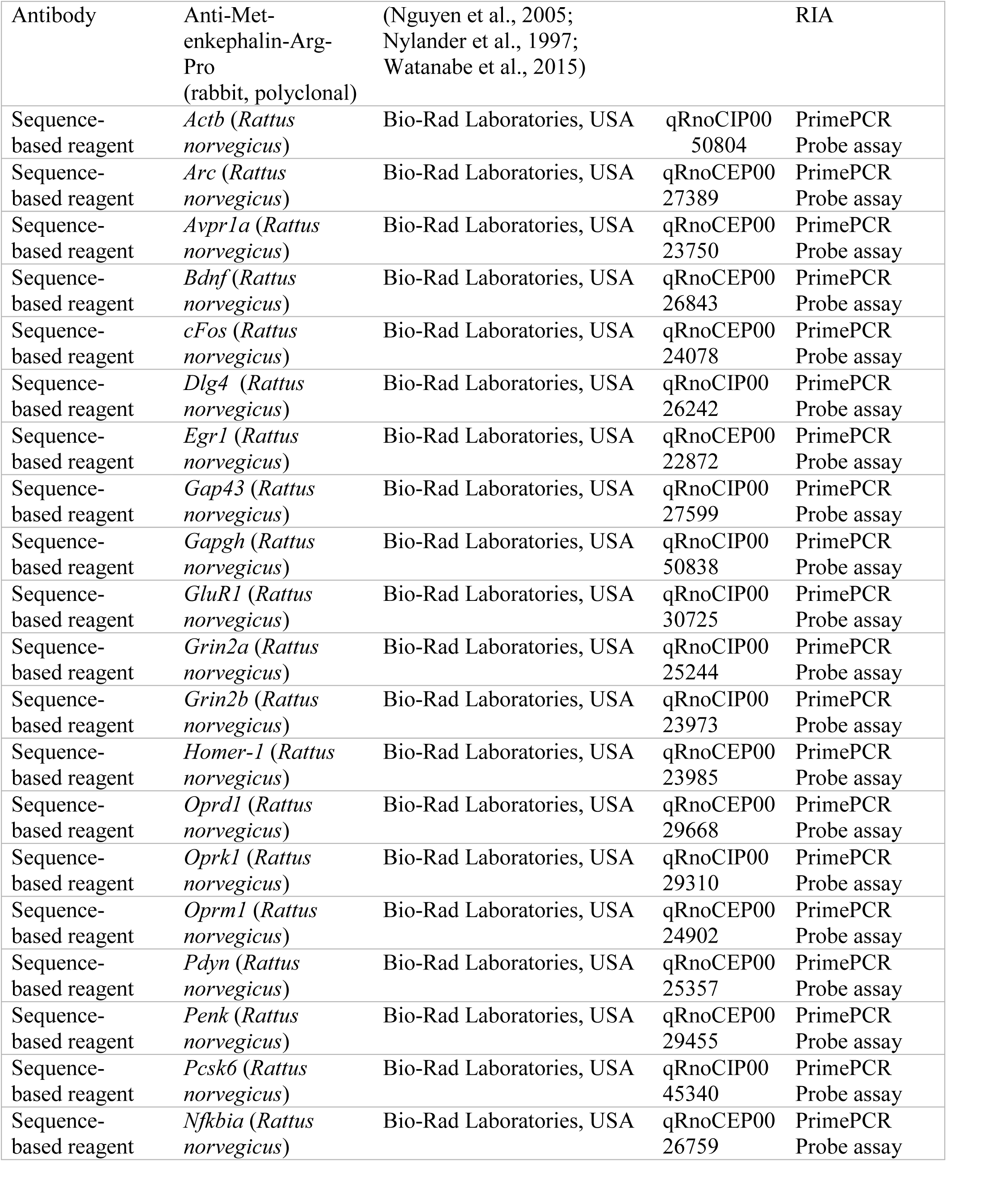
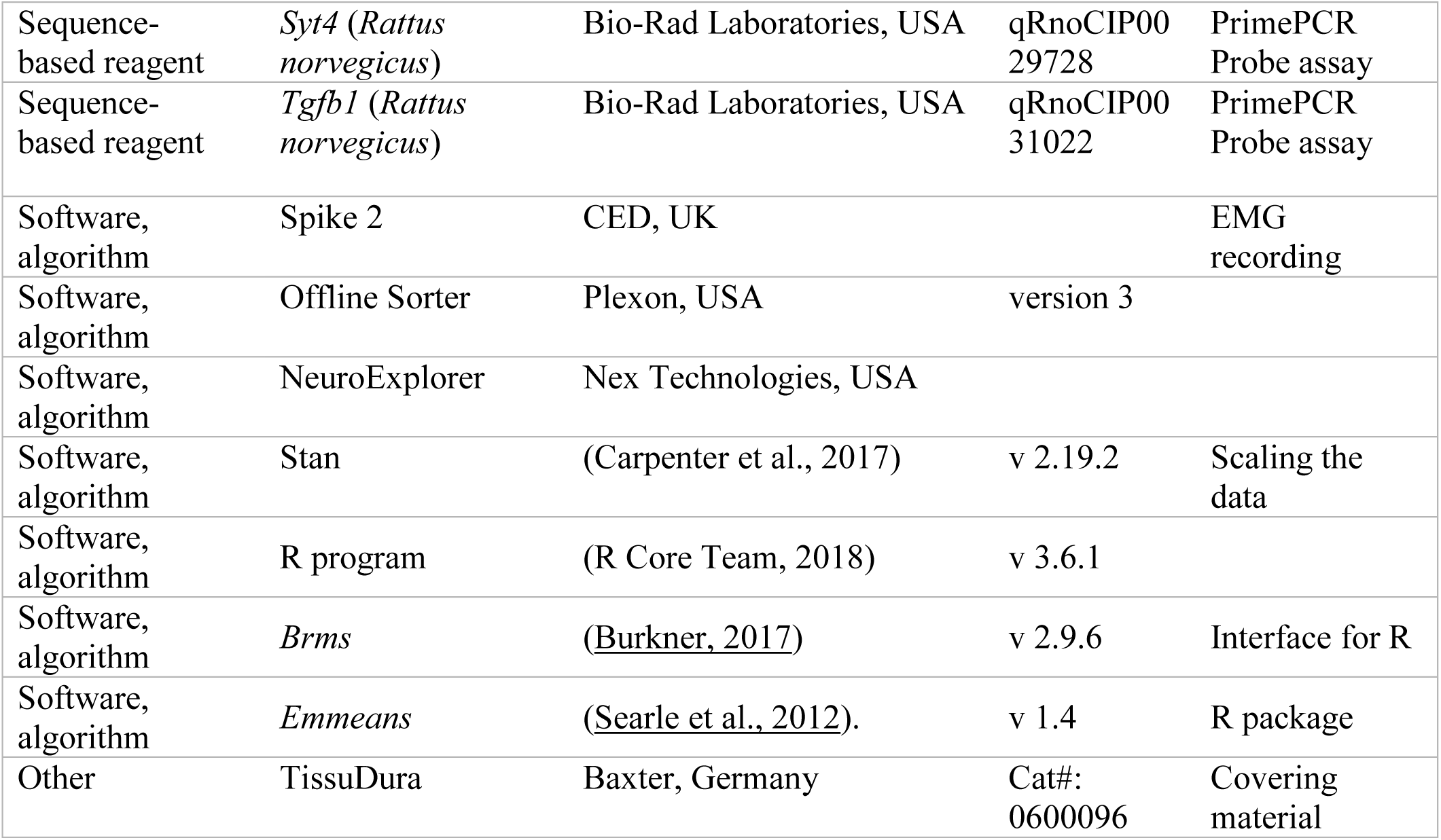

#### Animals

Male Wistar Hannover (Charles River Laboratories, Spain), 150-200 g body weight rats were used in behavioral, HL-PA and molecular experiments (***Figures 1, 3*** and ***4B-E***; ***Figure 1—figure supplements 1, 2, 3F-L***; ***Figure 3—figure supplements 2,3***; ***Figure 4B—figure supplements 1F-O,2***). Male Sprague Dawley rats were used for analysis of HL-PA (***Figure 1— figure supplement 3B-E***), in electrophysiological experiments (***Figure 2, Figure 2—figure supplements 1,2***) (Taconic, Denmark; 150-400 g body weight) and for hypophysectomy (***Figure 4A*** and ***Figure 4— figure supplement 1B-E***) (Charles River Laboratories, France; 115-125 g body weight). The animals received food and water ad libitum and were kept in a 12-h day-night cycle at a constant environmental temperature of 21°C (humidity: 65%). Approval for animal experiments was obtained from the Malmö/Lund ethical committee on animal experiments (No.: M7-16), and the ethical committee of the Faculty of Medicine of Porto University and Portuguese Direção-Geral de Alimentação e Veterinária (No. 0421/000/000/2018).

#### Surgery

The animals were anesthetized with sodium pentobarbital (I.P.; 60 mg/kg body weight, as an initial dose and then 6 mg/kg every hour). If needed, the anesthesia was reinforced with ≈1.5% isoflurane (IsoFlo, Abbott Laboratories, Norway) in a mixture of 65% nitrous oxide-35% oxygen. Core temperature of the animals was controlled using a feedback-regulated heating system. In the experiments involving electrophysiological recordings, the rats were ventilated artificially via a tracheal cannula and the expiratory CO_2_ and mean arterial blood pressure (65–140 mmHg) was monitored continuously in the right carotid artery.

##### Aspiration brain injury and spinal cord transection

Experimental design included either the UBI alone or the UBI which was preceded by a complete spinal cord transection. In the UBI-only experiments, anaesthetized rats were placed on a surgery platform with stereotaxic head holder. The rat head was fixed in a position in which the bregma and lambda were located at the same horizontal level. After local injection of lidocaine (Xylocaine, 3.5 mg/ml) with adrenaline (2.2 μg/ml), the scalp was cut open and a piece of the parietal bone located 0.5 – 4.0 mm posterior to the bregma and 1.8 – 3.8 lateral to the midline (Paxinos & Watson, 2007) was removed. The part of the cerebral cortex located below the opening that includes the hind-limb representation area of the sensorimotor cortex (HL-SMC) was aspirated with a metallic pipette (tip diameter 0.5 mm) connected to an electrical suction machine (Craft Duo-Vec Suction unit, Rocket Medical Plc, UK). Care was taken to avoid damaging the white matter below the cortex. After the ablation, bleeding was stopped with a piece of Spongostone and the bone opening was covered with a piece of TissuDura (Baxter, Germany). For sham operations, animals underwent the same anesthesia and surgical procedures, but the cortex was not ablated.

In the experiments in which UBI was preceded by the spinal cord transection, the anaesthetized animals were first placed on a surgery platform and the skin of the back was incised along the midline at the level of the superior thoracic vertebrae. After the back muscles were retracted to the sides, a laminectomy on the T2 and T3 vertebrae was performed. Then, the spinal cord between the two vertebrae was completely transected using a pair of fine scissors (Bakalkin & Kobylyansky, 1989). A piece of Spongostan (Medispon® (MDD sp. zo.o., Toruń, Poland) was placed between the rostral and caudal stumps of the spinal cord. The completeness of the transection was confirmed by (i) inspecting the cord during the operation to ensure that no spared fibers bridged the transection site and that the rostral and caudal stumps of the spinal cord are completely retracted; and (ii) examining the spinal cord in all animals after termination of the experiment. Following the spinalization procedures, the rats were moved to the stereotaxic frame and UBI was inflicted as described above. After completion of all surgical procedures, the wounds were closed by the 3-0 suture (AgnTho’s, Sweden) and the rat was kept under infrared radiation lamp to maintain body temperature during monitoring of postural asymmetry (up to 3 h) and during EMG recordings.

To verify the UBI site the rats were perfused with 4% paraformaldehyde and the brains were removed from the skulls. Following post fixation overnight in the same fixative, the brains were soaked in phosphate-buffered saline for 2 days, dissected into blocks and the blocks containing the lesion area were cut into 50 µm sections using a freezing microtome. Every fourth section was mounted on slides and stained for Nissl with modified Giemsa solution (Sigma-Aldridge, USA; 1:5 dilution). The left drawing in ***Figure 1A*** shows the location of the right hindlimb representation area on the rat brain surface (adapted from (Frost et al., 2013)).

##### Hypophysectomy

The hypophysectomy was performed at the Charles River Laboratories (France) and all the surgery-related procedures, including postoperative care and transportation of animals, were performed according to the ethical recommendations of that company. The procedure for transauricular hypophysectomy, performed under isoflurane anesthesia, was described elsewhere (Koyama, 1962). Briefly, a hypodermic needle fitted to a plastic syringe was introduced into the external acoustic meatus until its tip reached the medial wall of the tympanic cavity. The needle was then pushed slightly further, so that its tip pierced the bone and entered the pituitary capsule. The hypophysis was then sucked into the syringe. The success of hypophysectomy was assessed by visual inspection of the hypophyseal region of the skull under a microscope following animal sacrifice and removal of the brain. Only data obtained in rats in which complete hypophysectomy was confirmed were included in the analysis. Sham-operated rats underwent an identical procedure except that the needle was not introduced into the pituitary capsule. Following the hypophysectomy, the animals were given 3 weeks of recovery before initiating the UBI experiments.

#### Behavioral tests

Experiments were performed between 10:00 and 15:00 h over the course of the week preceding surgeries (pre-training) and one-day post-surgery (testing).

##### Beam-walking test (BWT) (Feeney et al., 1982)

The BWT apparatus consisted of a horizontally placed wooden beam, 130 cm long and 1.4 cm wide, which was elevated 55 cm above the surface of a table. One end of the beam was free, while another was connected to a square platform (10 x 10 cm, with the floor and two sidewalls painted in black) leading to the rat’s home cage. The rats were trained to walk along the beam from its free end to their home cage. Training continued during two consecutive days preceding the brain surgery, with three daily sessions and six trials per session. On the first, second and third trials, the rat was placed on the beam close to the platform, at the midpoint, and at the starting point (free end of the beam), respectively. The rat was considered trained if it performed the task within 80 seconds on the second training day. On the day following the UBI/Sham surgery, each rat was given one session consisting of three trials (runs). Each run was video-recorded for further offline analysis by an observer blind to the treatment groups. The number of times the left and the right hindlimbs slipped off the beam were registered and averaged across all runs of a given session.

##### Ladder Rung Walking Test (LRWT) (Metz & Whishaw, 2002)

The horizontal LRWT apparatus, 100 cm long × 20 cm high, consisted of sidewalls made of clear Plexiglas and metal grid floor. The width of the apparatus was approximately 12 cm. The floor was composed of removable stainless steel bars (rungs), 3 mm in diameter, spaced 1 cm apart (center-to-center). The ladder was placed 30 cm above the surface of a table and was connected to the animal’s home cage at one of its ends. Its opposite end was open and served as a starting point. On each trial, the rat was placed on the starting point and allowed to cross the ladder to enter the home cage. The width of the apparatus was adjusted to the size of the animal in order to prevent the animal from running in the reverse direction. During training (one session consisting of five trials), every second bar was removed, so that the rungs were spaced regularly at 2 cm intervals.

During post-surgery testing (one session of five trials), the rung spacing pattern was modified in order to increase the difficulty of the task. In particular, five distinct irregular spacing patterns were implemented in the testing session: 001101, 011010100, 1010011100, 1000011010 and 10001011000, where 1 denotes a rung and 0 a missing rung. However, the rung spacing patterns and the order of their presentation were the same for all rats. Each ladder run was video-recorded for further offline analysis by an observer blind to the surgery type. A total number of steps made by the left and right hindlimbs during each run and the number of times the limbs slipped between the rungs were registered. The limb slips/total steps ratio was averaged across the five testing trials and was used as an error score.

#### Analysis of postural asymmetry

The HL-PA was recorded as described previously (Bakalkin & Kobylyansky, 1989). Briefly, the postural asymmetry measurement was performed under pentobarbital (60 mg/kg, I.P.) anesthesia, or isoflurane anesthesia when rats exposed to unilateral brain injury were analyzed 1 or more days after the surgery. The level of anesthesia was characterized by a barely perceptible corneal reflex and a lack of overall muscle tone. The rat was placed in the prone position on the 1-mm grid paper, and the hindlimbs were straightened in the hip and knee joints by gently pulling them backwards for 5-10 mm to reach the same level Then, the limbs were set free and the magnitude of postural asymmetry was measured in millimeters as the length of the projection of the line connecting symmetric hindlimb distal points (digits 2-4) on the longitudinal axis of the rat. The procedure was repeated six times in immediate succession, and the mean HL-PA value for a given rat was calculated and used in statistical analyses. The measurements were performed 0.5, 1 and 3 hours after the brain injury, or at other time points as shown on figures. In a separate group of rats (***Figure 1—figure supplement 1C,D***), HL-PA was assessed 1, 4, 7 and 14 days after the UBI or sham surgery under the isoflurane anesthesia (1.5% isoflurane in a mixture of 65% nitrous oxide and 35% oxygen). The rat was regarded as asymmetric if the magnitude of HL-PA exceeded the 1 mm threshold (see statistical section). The limb displacing shorter projection was considered as flexed.

In a subset of the rats with UBI or sham surgery (n = 11 and 10, respectively), the hindlimbs were stretched by gently pulling two threads glued to the nails of the middle three toes of the both legs. In another subset (n = 6), the skin of the hindlimbs including and distal to the ankle joints was fully anesthetized by a topical application of 5% lidocaine cream 10 min before the assessments of HL-PA in rats with UBI. The absence of the pedal withdrawal reflexes following lidocaine application was confirmed in awake rats by pinching the skin between the toes with blunt forceps. None of these two procedures affected the resulting HL-PA suggesting that HL-PA formation does not dependent on tactile input from the hind paw.

Throughout the main text and the supplement, data are shown for the prone position, with the exception of ***Figure 1—figure supplement 2F-J***, which displays results for the supine position. For analysis in the supine position, the rat was placed in a V-shaped trough, a 90° - angled frame located on a leveled table surface with the 1-mm grid sheet; otherwise, the procedure was the same as for the prone position. The HL-PA values and the probability to develop asymmetry (P_A_) were essentially the same for both positions.

The postural asymmetry analysis was blind to the observer excluding the analysis combined with the EMG. The “reverse design” results shown on ***Figure 1F*** were replicated by two groups in different laboratories (***Figure 1—figure supplement 3B-E and F-L***, respectively).

#### EMG experiments

##### Electromyography recordings

Core temperature was maintained between 36 and 38°C using a thermostatically controlled, feedback-regulated heating system. The EMG activity of the extensor digitorum longus (EDL), interrossi (Int), peroneaus longus (PL) and semitendinosus (ST) muscles of both hindlimbs were recorded as described previously (Schouenborg et al., 1992; Weng & Schouenborg, 1996). EMG recordings were initiated approximately 3 hours after spinalization that was 2 h and 20 min after the UBI. Recordings were performed using gauge stainless steel disposable monopolar electrodes (DTM-1.00F, The Electrode Store, USA). The electrodes were insulated with Teflon except for ∼200 μm at the tip. The impedance of the electrodes was from 200 to 1000 kΩ. For EMG recordings, a small opening was made in the skin overlying the muscle, and the electrode was inserted into the mid-region of each muscle belly. A reference electrode was inserted subcutaneously in an adjacent skin region. The electrode position was checked by passing trains (100 Hz, 200 ms) of cathodal pulses (amplitude < 30 µA, duration 0.2 ms). The EMG signal was recorded with Spike 2 program (CED, UK) with a sampling rate of 5000 Hz. Low and high pass filter was set at 50000 Hz and 500 respectively. Generally, the EMG activity of three or four pairs of hind limb muscles was recorded simultaneously in each experiment / rat.

##### Cutaneous stimulation

Digits 2, 3, 4 and 5, and the heel were stimulated to induce reflex responses according to muscle’s receptive field as reported previously (Schouenborg et al., 1992; Weng & Schouenborg, 1996). Nociceptive stimulation was performed by intracutaneous cathodal electrical stimulation using the same electrodes as for EMG recording. The same type of electrodes was used as an anode and was placed subcutaneously in a skin flap well outside the stimulation area.

To detect the stimulation intensity that induce the maximal reflex in each muscle, graded current pulses (1 ms, 0.1 Hz) were used ranging mostly from 1 to 30 mA, occasionally up to 50 mA in anesthetized rats. The reflex threshold was defined as the lowest stimulation current intensity evoking a response at least in 3 out of 5 stimulations. If a muscle response was induced by stimulation at more than one site, the lowest current was taken as a threshold value. For EMG data collection, the current level that induced submaximal EMG responses from both legs, usually at 5 – 20 mA was chosen. This was usually 2 – 3 times higher than the threshold currents. The same current level was used on symmetrical points from the most sensitive area on both paws. For each site EMG responses from 18 – 20 stimulations at 0.1 Hz frequency were collected. No visible damage of the skin, or marked changes in response properties at the stimulation sites, were detectable at these intensities.

#### EMG data analysis

##### EMG amplitude

The spikes from Spike2 EMG data files were sorted with Offline Sorter (version 3, Plexon, USA). The EMG amplitude (spike number) from different muscles was calculated with NeuroExplorer (Nex Technologies, USA). To avoid stimulation artifacts, spikes from the first one or two stimulations were removed from further analysis. The number of spikes was calculated from 16 consecutive stimulations thereafter. The EMG thresholds and responses registered from 0.2 to 1.0 sec corresponding to C fiber evoked reflexes, were analyzed.

EMG activity was recorded in response to 18 stimulations with most responses stable between the stimulations (Supplement Fig. 4A,B). Peristimulus histograms denote that a main fraction of the responses was recorded in the 0.2 – 1.0 sec interval and were likely elicited through activation of C-fibers. Responses to the 2^nd^ to 17^th^ stimulations recorded between 0.2 – 1.0 sec were included in statistical analysis.

#### Effects of serum, neurohormones and antagonists of opioid (naloxone) and vasopressin V1B (SSR-149415) receptors on HL-PA development

Serum was collected from 3 animals in each group of rats with transected spinal cord 3 h after the UBI or sham surgery, kept at -80°C until use, and administered intravenously (0.3 mL / rat) to rats under pentobarbital anesthesia 10 min after complete spinal cord transection.

Serum and Arg-vasopressin were administered into the cisterna magna (intracisternal route; 5 microliters/rat) (Ramos et al., 2019; Xavier et al., 2018) of intact rats under pentobarbital anesthesia, which was followed by the spinal cord transection 10 min later. HL-PA was analyzed at the 0.5, 1 and 3 h time points after injection.

SSR-149415 (5 mg/ml/kg, n=12) dissolved in a mixture of DMSO (10%) and saline (90%), or vehicle alone (n = 8) was administered I.P. 10 min before spinal cord transection. This was followed by either the left-side UBI (SSR-149415: n = 6; vehicle: n = 5) or intravenous administration of serum from UBI rats (SSR-149415: n = 6; vehicle: n = 3). In rodents, effects of SSR149415 develop within 0.5-1 h and last for 4-6 h after administration (Ramos et al., 2006; Serradeil-Le Gal et al., 2002). Naloxone (5 mg/ml/kg in saline) or saline alone was injected I.P. 2 h after delivering the UBI (naloxone: n = 6; saline: n = 6) or after injecting the UBI serum (naloxone: n = 6; saline: n = 3) to rats with transected spinal cord.

#### Analysis of mRNA levels by quantitative RT-PCR (qRT-PCR)

Total RNA was purified by using RNasy Plus Mini kit (Qiagen, Valencia, CA, USA). RNA concentrations were measured with Nanodrop (Nanodrop Technologies, Wilmington, DE, USA). RNA (1 μg) was reverse-transcribed to cDNA with the iScript ™ cDNA Synthesis Kit (Bio-Rad Laboratories, CA, USA) according to manufacturer’s protocol. cDNA samples were aliquoted and stored at -20°C. Assay was described elsewhere (Kononenko et al., 2017; Kononenko et al., 2018). cDNAs were mixed with PrimePCR™ Probe assay (Table S2) and iTaq Universal Probes supermix (Bio-Rad) for qPCR with a CFX384 Touch™ Real-Time PCR Detection System (Bio-Rad Laboratories, CA, USA) according to manufacturer’s instructions. mRNA levels of genes of interest were normalized to geometric mean of expression levels of two control genes *Actb* and *Gapdh* selected by geNORM program (https://genorm.cmgg.be/ and (Kononenko et al., 2017; Kononenko et al., 2018; Vandesompele et al., 2002)). In each experiment, internal control gene-stability measure M (Vandesompele et al., 2002) did not exceed the established limit of 0.5.

#### Radioimmunoassay (RIA)

The procedure was described elsewhere (Christensson-Nylander et al., 1985; Merg et al., 2006). Briefly, 1 M hot acetic acid was added to finely powdered frozen tissues, and samples were boiled for 5 min, ultrasonicated and centrifuged. Tissue extract was run through SP-Sephadex ion exchange C-25 column, and peptides were eluted and analyzed by RIA. Anti-Dynorphin B antiserum showed 100% molar cross-reactivity with big dynorphin, 0.8% molar cross-reactivity with Leu-morphine (29 amino acid C-terminally extended Dynorphin B), and <0.1% molar crossreactivity with Dynorphin A (1–17), Dynorphin A (1–8), α-neoendorphin, and Leu-enkephalin (Yakovleva et al., 2006). Cross-reactivity of Leu-enkephalin-Arg antiserum with Dynorphin B and Leu- and Met-enkephalin was <0.1% molar, with α-neoendorphin 0.5% molar, with Dynorphin A (1–8) 0.7% molar, with Met-enkephalin-Arg-Phe 1% molar and with Met-enkephalin-Arg 10% molar. Cross-reactivity of Met-enkephalin-Arg-Phe antiserum with Met-enkephalin, Met-enkephalin-Arg, Met-enkephalin-Arg-Gly-Leu, Leu-enkephalin and Leu-enkephalin-Arg was <0.1% molar (Nylander et al., 1997). Our variant of RIA readily detected Dynorphin B and Leu-enkephalin-Arg in wild-type mice (Nguyen et al., 2005) but not in *Pdyn* knockout mice; thus the assay was highly specific and not sensitive to the presence of contaminants. The peptide content is presented in fmol/mg tissue.

#### Statistical Analysis

##### Postural asymmetry and NWR

Predictors and outcomes were centered and scaled before Bayesian Regression Models were fitted by calling Stan 2.19.2 (Carpenter et al., 2017) from *R* 3.6.1 (R Core Team, 2018) using *brms* 2.9.6 (Burkner, 2017) interface. To reduce the influence of outliers, models used Student’s *t* response distribution with identity link function. Models had no intercepts with indexing approach to predictors (McElreath, 2019). Default priors were provided by *brms* according to Stan recommendations (Gelman, 2019). Residual SD and group-level SD were given weakly informative prior student_t(3, 0, 10). Additional parameter ? of Student’s distribution representing the degrees of freedom was given wide gamma prior gamma(2, 0.1). Group-level effects were given generic weakly informative prior normal(0, 1). Removal of non-significant confounders was attempted in stepwise fashion comparing models by exact 10-fold cross-validation. Four Markov chain Monte Carlo (MCMC) chains of 40000 iterations were simulated for each model, with a warm-up of 20000 runs to ensure that effective sample size for each estimated parameter exceeded 10000 (Kruschke, 2015) producing stable estimates of 95% highest posterior density continuous intervals (HPDCI). MCMC diagnostics were performed according to Stan manual.

Median of the posterior distribution, 95% HPDCI and adjusted P-values are reported as computed by R package *emmeans* 1.4 (Searle et al., 2012). Adjusted P-values were produced by frequentist summary in *emmeans* using the multivariate t distribution with the same covariance structure as the estimates. The asymmetry and contrast between groups were defined as significant if the corresponding 95% HPDCI interval did not include zero value and, simultaneously, adjusted P-value was ≤ 0.05. R scripts are available upon request.

The 99^th^ percentile of the HL-PA magnitude in rats after sham surgery (n = 36) combined at the 2 or 3 h time points was 1.1 mm. Therefore, the rats with HL-PA magnitude > 1 mm threshold were defined as asymmetric. The probability P_A_ to develop HL-PA above 1 mm in magnitude was modelled with Bernoulli response distribution and logit link function. The UBI effects remained significant for models with thresholds of 2 or 3 mm.

In the EMG analysis, asymmetry in stimulation threshold (Thr) and a spike number (SN) in 0.2 – 1 sec EMG responses for each pair of hindlimb muscles analyzed was assessed using the Contra- /Ipsilesional asymmetry indexes AI_Thr_ (AI_Thr_ = log_2_ [Thr_contra_ / Thr_ipsi_]) and AI_SN_ (AI_SN_ = log_2_ [(1+SN_contra_)/(1+SN_ipsi_)]), where *contra* and *ipsi* designate the Contralateral and Ipsilesional sides relative to the brain injury side. *Operation type* (UBI *vs*. sham) was the factor of interest analyzed for each muscle (EDL stimulated at D3, D4 and D5; Int at D2, D3, D4 and D5; PL at D4 and D5; and ST at the heel). Data recorded at stimulation of more than one site were processed as replicates for a given muscle. The number of rats for each pair of muscles in each UBI and sham group is shown in ***Figure 2—figure supplement 2***.

The AI_SN_ was fitted using linear multilevel model with fixed effects of *muscle* (EDL, Int, PL and ST) interacting with *operation type* (UBI vs. sham) and log-transformed *recoding current*. The *sweep’s number* was a fixed effect confounder with non-significant effect, showing that the AI_SN_ was not significantly affected by wind-up. The AI_Thr_ was fitted using the similar linear multilevel model without the *recording current* and *sweep’s number* factors. To get rid of No-U-Turn Sampler warnings, parameters adapt_delta=0.999 and max_treedepth=13 were used.

##### Gene expression and opioid peptide analyses

First, the mRNA levels of 20 neuroplasticity (*Arc, Bdnf, cFos, Dlg4, Egr1, Homer-1, Gap43, GluR1, Grin2a, Grin2b, Nfkbia, Pcsk6, Syt4* and *Tgfb1*), and opioid and vasopressin (*Penk, Pdyn, Oprm1, Oprd1, Oprk1* and *Avpr1a*) genes, and the levels of opioid peptides dynorphin B, Leu-enkephalin-Arg and Met-enkephalin-Arg-Pro were compared separately for the left and right halves of the lumbar spinal cord between the left UBI (n = 12) and left sham (n = 11) rat groups. Only *Avpr1a* out of four genes of the vasopressin system (*Avpr1a, Avpr1b, Avpr2, and Avp)* was found to be expressed in the lumbar spinal cord and therefore included in the statistical analyses. The mRNA and peptide levels were compared between the animal groups for the left and right spinal domains separately using Mann-Whitney test followed by Bonferroni correction for a number of tests (correction factor for mRNAs was 40, for peptides 6). Results were considered significant if P value corrected for multiple comparisons (P_adjusted_) did not exceed 0.05. Log fold change (logFC) was computed as a difference of median log_2_-scaled expression values.

Second, the expression asymmetry index (eAI) defined as log_2_-scaled ratio of expression levels in Contra and Ipsilesional spinal domains (log_2_[Contra/Ipsi]), was compared between the rat groups. Comparison of eAI was performed using Mann-Whitney test followed by Bonferroni correction for multiple tests (correction factor was 20).

Heatmaps of expression levels and eAI were constructed using data (0,1)-standardized for each gene by subtraction of the median value and division by an inter-quartile range. In analysis of expression levels, standardization was applied to log_2_-scaled expression levels pooled for the left and the right spinal domains.

##### Intra- and inter-regional gene coexpression patterns

Spearman’s rank correlation coefficient was calculated for all gene pairs in each area (N = 190) and between the areas (N = 400). To circumvent effects of differences in a number of animals between the groups (caused by differences in the number of rats or by missing values) on outcome of statistical analyses, the following procedure was applied. For a given pair of genes, animals with missing expression levels were excluded from calculations. For the group with smaller number of remaining animals (let *N* denote this number) correlation coefficient was calculated in a standard way. For the other group correlation coefficient was calculated for all subsets consisting of *N* animals, and the median was taken. The procedure was applied separately for each pair of genes in each analysis. Significance of correlation coefficients was assessed using pspearman R package with precomputed null distribution (i.e., *approximation* parameter set to “*exact*”).

Statistical comparison of gene-gene coordination strength between the animal groups was performed by applying Wilcoxon signed-rank test to the set of absolute values of all correlation coefficients and, separately, to the set of absolute values of significant correlation coefficients (i.e., having associated P-value not exceeding 0.05 for at least one animal group). As the comparison of gene-gene coordination strength ignored correlation signs, a separate analysis was performed to assess differences in the proportion of positive and negative correlations between animal groups. This assessment was performed separately for the sets of (i) all and (ii) significant correlation coefficients using the Fisher’s Exact test with 2×2 contingency table.

## Acknowledgements

This paper is dedicated to Professors Boris I. Klement’ev and Genrich A. Vartanian for their outstanding postural asymmetry studies, which are mostly unknown to the Western scientific community and which set the stage for the present investigation. We are grateful to Dr. Michael Ossipov for discussion and manuscript processing, and Dr. Aleh Yahorau for help with biochemical assays and figure preparation. The study was supported by the Swedish Science Research Council (Grants K2014-62X-12190-19-5 and 2019-01771-3), P.O. Zetterling foundation, Uppsala University, and grants of the Government of the Russian Federation (14.W03.31.0031) and the Russian Scientific Foundation (17-14-01338).

## Additional information

### Funding

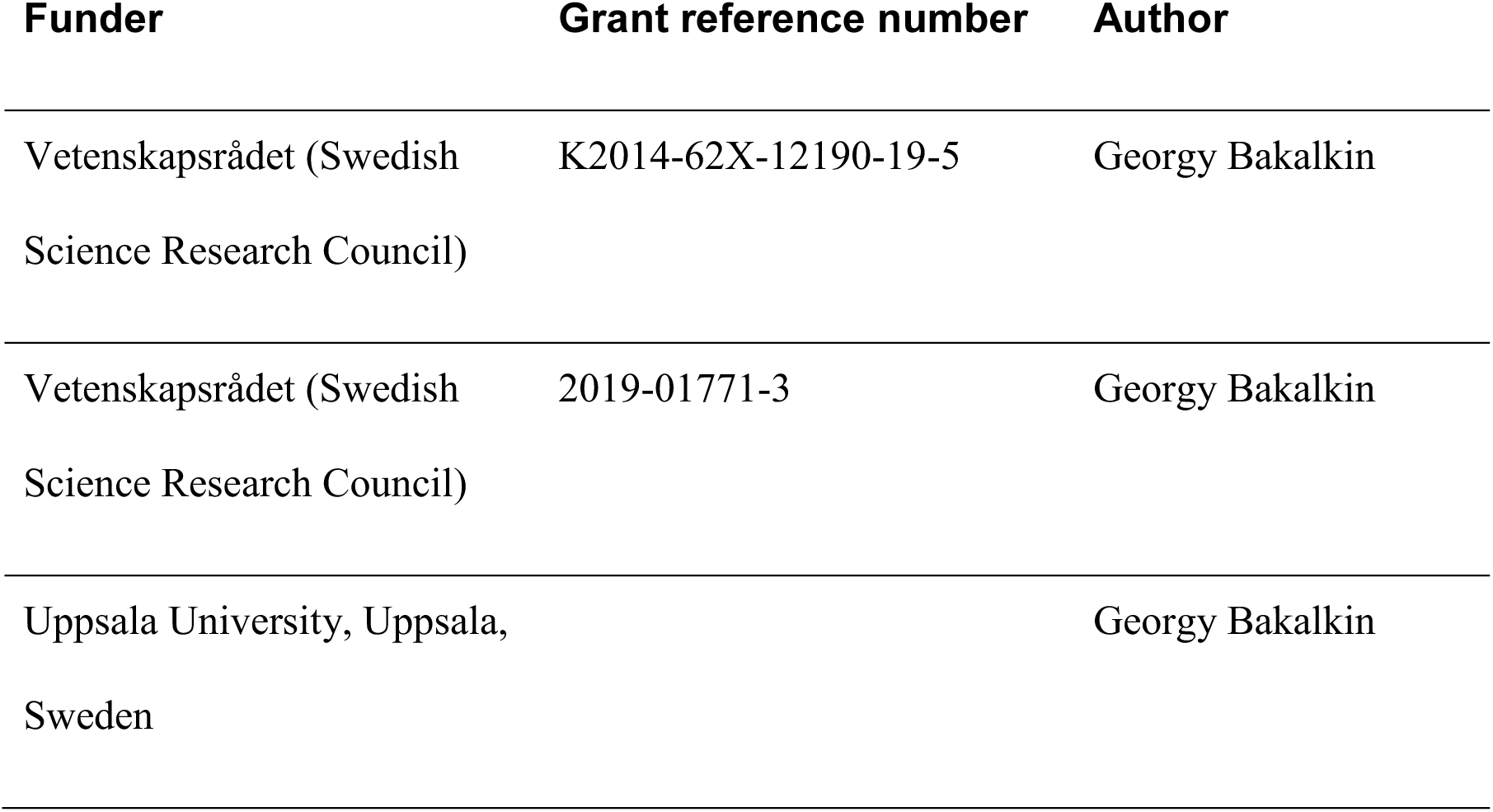
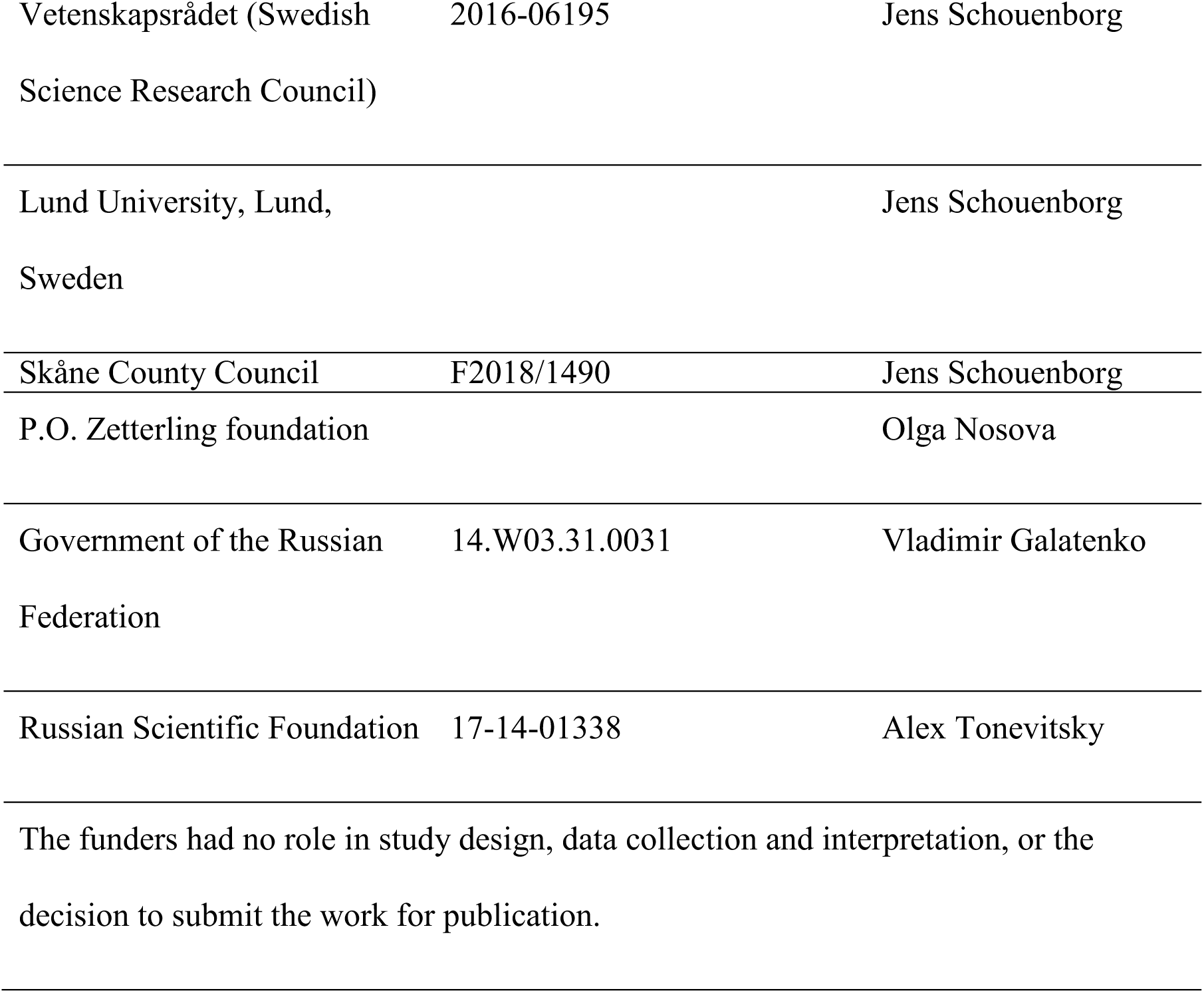

### Author contributions

Nikolay Lukoyanov, Formal analysis, Investigation, Conceptualization, Supervision, Methodology, Writing – original draft, Writing – review and editing; Hiroyuki Watanabe, Data curation, Formal analysis, Investigation, Methodology; Liliana S. Carvalho, Data curation, Formal analysis, Investigation, Methodology; Olga Nosova, Data curation, Formal analysis, Investigation,Methodology; Daniil Sarkisyan, Formal analysis, Investigation, Methodology, Writing – original draft; Mengliang Zhang, Data curation, Formal analysis, Investigation, Methodology, Writing – review and editing; Marlene Storm Andersen, Data curation, Formal analysis, Investigation; Elena A. Lukoyanova, Data curation, Formal analysis, Investigation; Vladimir Galatenko, Formal analysis, Investigation, Methodology, Writing – original draft preparation; Alex Tonevitsky, Formal analysis, Investigation, Resources; Igor Bazov, Data curation, Formal analysis, Investigation; Tatiana Iakovleva, Data curation, Formal analysis, Investigation; Jens Schouenborg, Conceptualization, Resources, Supervision, Writing – review and editing; Georgy Bakalkin, Conceptualization, Resources, Supervision, Funding acquisition, Project administration, Writing – original draft preparation; Writing – review and editing.

### Ethics

Animal experimentation: The animals received food and water ad libitum and were kept in a 12-h day-night cycle at a constant environmental temperature of 21°C (humidity: 65%). Approval for animal experiments was obtained from the Malmö/Lund ethical committee on animal experiments (No.: M7-16), and the ethical committee of the Faculty of Medicine of Porto University and Portuguese Direção-Geral de Alimentação e Veterinária (No. 0421/000/000/2018).

## Additional files

### Supplementary files

Transparent reporting form.

### Data availability

All data generated or analyzed during this study are summarized in the manuscript, figures and supplementary files.

Source data files generated or analyzed during this study are included for Figures 1-4.

